# Coordinated poleward flux of sister kinetochore fibers drives chromosome alignment

**DOI:** 10.1101/2020.12.30.424837

**Authors:** Patrik Risteski, Domagoj Božan, Mihaela Jagrić, Agneza Bosilj, Nenad Pavin, Iva M. Tolić

## Abstract

Chromosome alignment at the spindle equator promotes proper chromosome segregation and depends on pulling forces exerted at kinetochore fiber tips together with polar ejection forces. However, kinetochore fibers are also subjected to forces driving their poleward flux. Here we introduce a flux-driven centering model that relies on flux generated by forces within the overlaps of bridging and kinetochore fibers. This centering mechanism works so that the longer kinetochore fiber fluxes faster than the shorter one, moving the kinetochores towards the center. We developed speckle microscopy in human spindles and confirmed the key prediction that kinetochore fiber flux is length-dependent. Kinetochores are better centered when overlaps are shorter and the kinetochore fiber flux markedly slower than the bridging fiber flux. We identify Kif18A and Kif4A as overlap and flux regulators and NuMA as a fiber coupler. Thus, length-dependent sliding forces exerted by the bridging fiber onto kinetochore fibers promote chromosome alignment.

## INTRODUCTION

Chromosome alignment at the spindle equator in metaphase is a hallmark of mitosis and is important for proper completion of mitosis (Fonseca et al., 2019; Maiato et al., 2017). Chromosome movements on the spindle that lead to their alignment are driven by pulling forces exerted by kinetochore microtubules that pull the kinetochores poleward and polar ejection forces exerted by non-kinetochore microtubules that push the chromosome arms away from the pole (Rieder and Salmon, 1994). The role of these forces in chromosome movements and alignment were explored in theoretical studies (Joglekar and Hunt, 2002; Civelekoglu-Scholey et al., 2006, 2013; Armond et al., 2015). The main mechanism of chromosome alignment in these models relies on polar ejection forces, which have a centering effect on chromosomes because these forces decrease away from the spindle pole (Ke et al., 2009).

Similarly to the polar ejection forces, pulling force generated by kinetochore microtubules can have a centering effect on chromosomes even though forces generated at the microtubule plus end do not depend on microtubule length. The centering effect arises due to motor proteins such as kinesin-8, which ‘measure’ microtubule length by binding along the microtubule lattice and walking all the way to the microtubule plus end, where they make microtubule dynamics length-dependent (Varga et al., 2006). Indeed, kinesin-8 is required for chromosome alignment at the spindle center (Mayr et al., 2007; Stumpff et al., 2008, 2012; West et al., 2002). Theoretical studies have shown that length-dependent microtubule catastrophe induced by kinesins or length-depending pulling forces can center kinetochores in yeast cells (Gardner et al., 2008; Mary et al., 2015; Gergely et al., 2016; Klemm et al., 2018). Thus, in addition to polar ejection forces, measuring of microtubule length by kinesins has an important contribution to chromosome centering.

However, this is not a complete picture of the forces that act on chromosomes. Kinetochore fibers (k-fibers) are also subjected to forces that drive their poleward flux (Forer et al., 1965; Hamaguchi et al., 1987; Hiramoto and Izutsu, 1977; Mitchison, 1989). This movement can be imagined as a conveyor belt-like transport where the whole k-fiber is being shifted towards the pole, while its minus ends depolymerize and plus ends polymerize. This complex process is driven and regulated by multiple motor proteins (Miyamoto et al., 2004; Ganem et al., 2005; Rogers et al., 2004; Steblyanko et al., 2020). It has been proposed that poleward flux of k-fibers is generated by motor-driven sliding of k-fibers with respect to interpolar microtubules (Mitchison, 2005), inspired by electron microscopy images of *Xenopus* extract spindles (Ohi et al., 2003). The mechanical interaction between k-fibers and the associated interpolar bundles called bridging fibers has been demonstrated by laser cutting of these fibers in human cells (Kajtez et al., 2016). Kinesin-5 activity contributes to the poleward flux of k-fibers and interpolar microtubules in *Drosophila* syncytial embryo mitosis (Brust-Mascher et al., 2009). How poleward flux of interpolar microtubules transmitted to k-fibers regulates forces acting on kinetochores has been explored in a theoretical model, which suggests that flux promotes tension uniformity on kinetochores, in agreement with experiments showing large variability in kinetochore tension in cells with abolished flux (Matos et al., 2009). Interestingly, physical coupling between k-fibers and the associated interpolar bundles (bridging fibers) is important not only for tension but also for chromosome alignment, given that optogenetic perturbation of bridging fibers led to chromosome misalignment (Jagrić et al., 2021). In these experiments, chromosome misalignment was accompanied by elongation of bridging microtubule overlaps. These recent findings, together with the idea that poleward flux is generated within bridging fibers and transmitted to k-fibers, open an interesting possibility that chromosome alignment, bridging microtubule overlaps and poleward flux are mutually related. Thus, the mechanism of chromosome alignment on the spindle is incompletely understood.

Here we hypothesize that poleward flux drives chromosome centering. We introduce a flux-driven centering model that relies on the interaction between bridging and k-fibers. The model describes a centering mechanism based on length-dependent pulling forces exerted by k-fibers onto the kinetochores. These forces increase with the overlap length between bridging and k-fibers and with the velocity difference between the fibers. To test this model, we developed a speckle microscopy assay on spindles of human cells, which allowed us to measure the flux of individual bridging and kinetochore microtubules. We found that at displaced kinetochores, the longer k-fiber undergoes flux at a higher velocity than the shorter one, which is at the core of the flux-driven centering because in this mechanism the faster flux of the longer k-fiber pulls the kinetochores in the direction of this fiber, i.e., towards the spindle center. Our experiments in which we performed a set of depletions of spindle proteins, together with theory, indicate that kinetochores are better centered when the overlaps between bridging and k-fibers are shorter and the kinetochore fiber flux markedly slower than the bridging fiber flux. Forces from the bridging fiber are transmitted to the k-fiber in a manner dependent on the coupling between bridging and k-fibers. We show that k-fibers flux slower after depletion of NuMA, indicating that NuMA couples the fibers, whereas k-fibers flux faster after depletion of Kif18A and/or Kif4A, which results in longer overlaps implying stronger coupling. Our results suggest that lateral length-dependent sliding forces that the bridging fiber exerts onto k-fibers promote the movement of kinetochores towards the spindle center.

## RESULTS

### Physical model for chromosome centering based on microtubule poleward flux

To explore the idea that microtubule poleward flux promotes kinetochore centering, we introduce a "flux-driven centering" model in which k-fibers laterally interact with bridging microtubules (**Fig. 1 A**). The central idea of our theory is that kinetochores are centered by pulling forces proportional to the overlaps of k-fibers and bridging microtubules. These forces are generated within the overlaps by the activity of motor proteins and by passive crosslinkers. When kinetochores are off-centered, the difference in the length of sister k-fibers leads to a difference in the length of antiparallel overlaps and thus of accumulated motors on either side, generating a centering force on the kinetochores (white arrows in **Fig. 1 A**). Similarly, a difference in the length of parallel overlaps and the number of accumulated crosslinkers also leads to centering of kinetochores (grey arrows in **Fig. 1 A**). Thus, the kinetochores become centered through tug-of-war between sister k-fibers, which is different from the previously proposed centering mechanism based on dynamics of k-fiber plus ends and polar ejection forces. By developing a theory for flux-driven centering, we explore how poleward flux centers kinetochores, and what aspects of the spindle are crucial for efficient centering.

**Figure 1.**
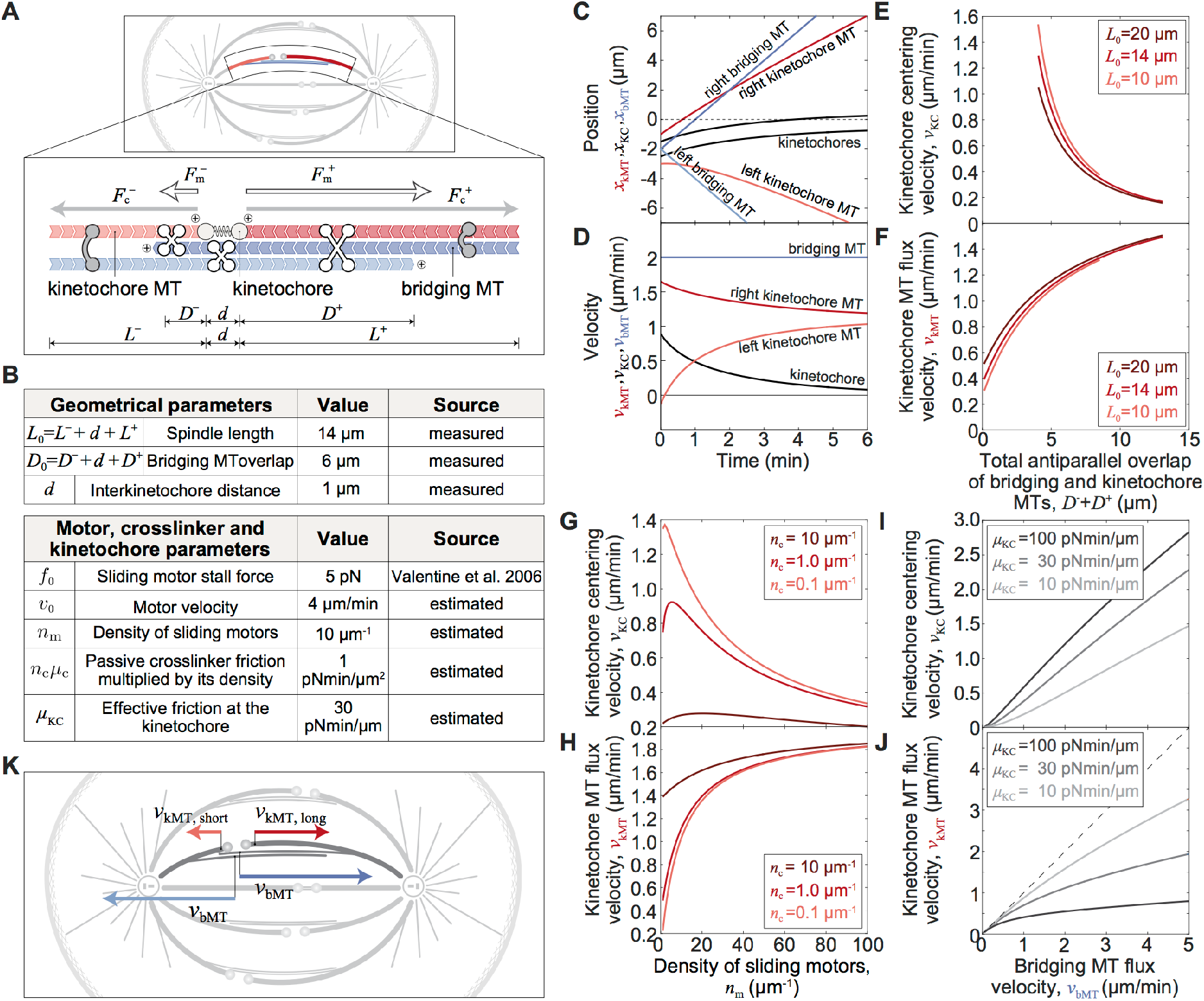
Theoretical model for chromosome alignment. (**A**) Scheme of mitotic spindle (top) and the scheme of the model (bottom). Kinetochore microtubules (red) extend from the edges toward elastically connected kinetochores (spring connecting circles). Bridging microtubules (blue) extend from the edges towards each other. Motor proteins (white X-shapes) exert forces, 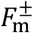, between antiparallel microtubules and passive crosslinkers (gray C-shapes) exert forces, 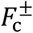, between parallel microtubules, where superscripts + and − denote the right and left sides, respectively. (**B**) Parameters of the model. Solution of the model showing time course of positions (**C**) and velocities (**D**) of kinetochores (black), kinetochore microtubules (red) and bridging microtubules (blue) for kinetochores initially displaced 2 µm. Kinetochore centering velocities and kinetochore microtubule flux velocities for different values of: (**E**) and (**F**) the length of antiparallel overlap and 3 values of spindle length, (**G**) and (**H**) sliding motor density and 3 values of passive crosslinker density, (**I**) and (**J**) bridging microtubule flux velocity and 3 values of effective friction at the kinetochore. Dashed line in (**J**) denotes the case in which bridging and kinetochore microtubule flux velocities are equal. (**K**) Scheme of mitotic spindle with flux velocities of kinetochore microtubules (red arrows) and bridging microtubules (blue arrows). Parameters for all panels are given in (**B**) if not stated otherwise.

A unique feature of our physical model is that motor proteins accumulate in the antiparallel overlaps between k-fibers and bridging microtubules, where they slide the microtubules apart. These sliding forces,

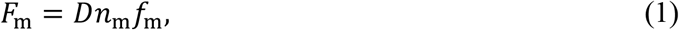

are proportional to the overlap length, *D*, based on *in vitro* experiments (Shimamoto et al., 2015). The force is also proportional to the linear density of motors, *n*_m_, each producing a force *f*_m_. Parallel overlaps between k-fibers and bridging microtubules are linked by passive crosslinkers, which help to transmit the sliding forces from the bridging microtubules to the k-fibers. Similar to the motor forces, the forces exerted by passive crosslinkers,

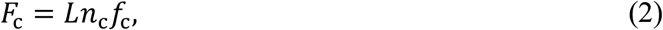

are proportional to the length of parallel overlaps, *L*, the linear density of crosslinkers, *n*_c_, and the force exerted by a single crosslinker, *f*_c_. The forces generated by motors on a k-fiber are opposed by the force exerted at the kinetochore, *F*_KC,_, and by passive crosslinkers,

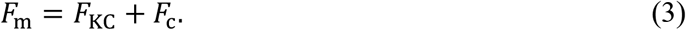

These are the main equations of the model, whereas a complete theory that includes not only the forces on k-fibers but also on bridging fibers, together with a force-velocity relationship for individual motors, and friction forces exerted by passive crosslinkers and kinetochores, is given in the Methods section.

To explore the key features of the centering mechanism, we displace the kinetochores in the model by 2 µm away from the spindle center and explore how they return to the center (**Methods**). The kinetochores approach the spindle center in several minutes for parameters typical for spindles in human (**Fig. 1 B, C**). When the kinetochores are displaced, the shorter k-fiber undergoes poleward flux at a slower velocity than the longer k-fiber, which is responsible for the movement of the kinetochores towards the spindle center (**Fig. 1 D**). The kinetochore centering velocity, which is equal to the half of the difference in poleward flux between two k-fibers, decreases as the kinetochores approach the center. The flux of both k-fibers is slower than the flux of bridging microtubules (**Fig. 1 D**), making the centering mechanism work by allowing the k-fibers to slide at different velocities.

To study what features of the system are crucial for efficient centering, we test the dependence of the centering velocity on the model geometry and the concentrations of motors and passive crosslinkers (**Fig. 1 E-H; Fig. S1**). We find that the length of antiparallel overlaps between bridging and k-fibers is the key determinant of the centering efficiency (**Fig. 1 E**). As the total antiparallel overlap between bridging and k-fibers increases from 4 µm to 13 µm, the centering velocity decreases roughly 7-fold (**Fig. 1 E**). For the same increase of overlap length, the k-fiber flux velocity increases and consequently the difference between k-fiber and bridging fiber velocities decreases from 1.0 µm/min to 0.5 µm/min (**Fig. 1 F**). Centering is better for short overlaps because the relative difference in the number of motors on either side is larger, resulting in a greater centering velocity. Similarly, centering velocity decreases with decreasing spindle length, but the effect is smaller than for the overlap length (**Fig. 1 E, F**).

By varying the density of motor proteins we find that the centering velocity has a maximum value below 20 motors per micron, around which the centering mechanism behaves optimally (**Fig. 1 G**). When the number of motors decreases from the optimum, the contribution of passive crosslinkers becomes larger than that of motors. This leads to worse centering because passive crosslinkers generate smaller centering forces than motors. When the number of motors increases from the optimal one, the centering becomes worse for a different reason. Here, the k-fiber flux velocity increases (**Fig. 1 H**) and thus the difference between k-fiber and bridging fiber velocities decreases, because a high number of motors slide k-fibers poleward at a high velocity, leading to slower centering. Additionally, when the density of passive crosslinkers increases, k-fibers slide poleward at higher velocities and consequently centering is slower (**Fig. 1 G, H**).

To explore the influence of the bridging fiber flux velocity on centering, we varied the velocity of motors in the absence of load, which is equal to the sliding velocity of the oppositely oriented bridging microtubules with respect to one another, or, in other words, twice the bridging microtubules flux velocity. The kinetochore centering velocity and the k-fiber flux velocity increase with the bridging microtubules flux velocity (**Fig. 1 I, J**). We also explored the influence of the parameter describing friction at the kinetochore on centering and found that kinetochores center faster and k-fiber flux decreases for larger values of this parameter. In all cases the k-fiber flux is slower than the bridging fiber flux, and this difference is larger when the bridging fiber flux is faster (**Fig. 1 J**).

Taken together, the flux-driven centering model provides a crucial prediction that is unique to this model: at displaced kinetochores, the longer k-fiber undergoes flux at a higher velocity than the shorter one (**Fig. 1 K; Movie 1**). The faster flux of the longer k-fiber pulls the kinetochores in the direction of this fiber, i.e., towards the spindle center. Thus, the difference in the flux of the sister k-fibers is the core of the centering mechanism.

### Speckle microscopy assay to follow the movement of individual microtubules within the spindle

To test the predictions of the flux-driven centering model experimentally, it is important to measure the poleward flux of different classes of microtubules (kinetochore and bridging), which requires analysis of the movements of individual microtubules. Flux is typically studied by using tubulin photoactivation (Mitchison, 1989), a method in which all the microtubules within the illuminated region are photoactivated, thus the movements of kinetochore and non-kinetochore microtubules cannot be distinguished. To overcome this issue, we developed an assay based on speckle microscopy (Waterman-Storer et al., 1998) to study microtubules within spindles of the human non-cancer immortalized epithelial cell line hTERT-RPE1 (from here on referred to as RPE1). By using a very low concentration (1 nM) of SiR-tubulin (Lukinavičius et al., 2014), we obtained speckled signal of SiR-tubulin in the spindle (**Fig. 2 A; Movie 2**), which comes from a few dye molecules within a resolution-limited region (Waterman-Storer and Salmon, 1998).

**Figure 2.**
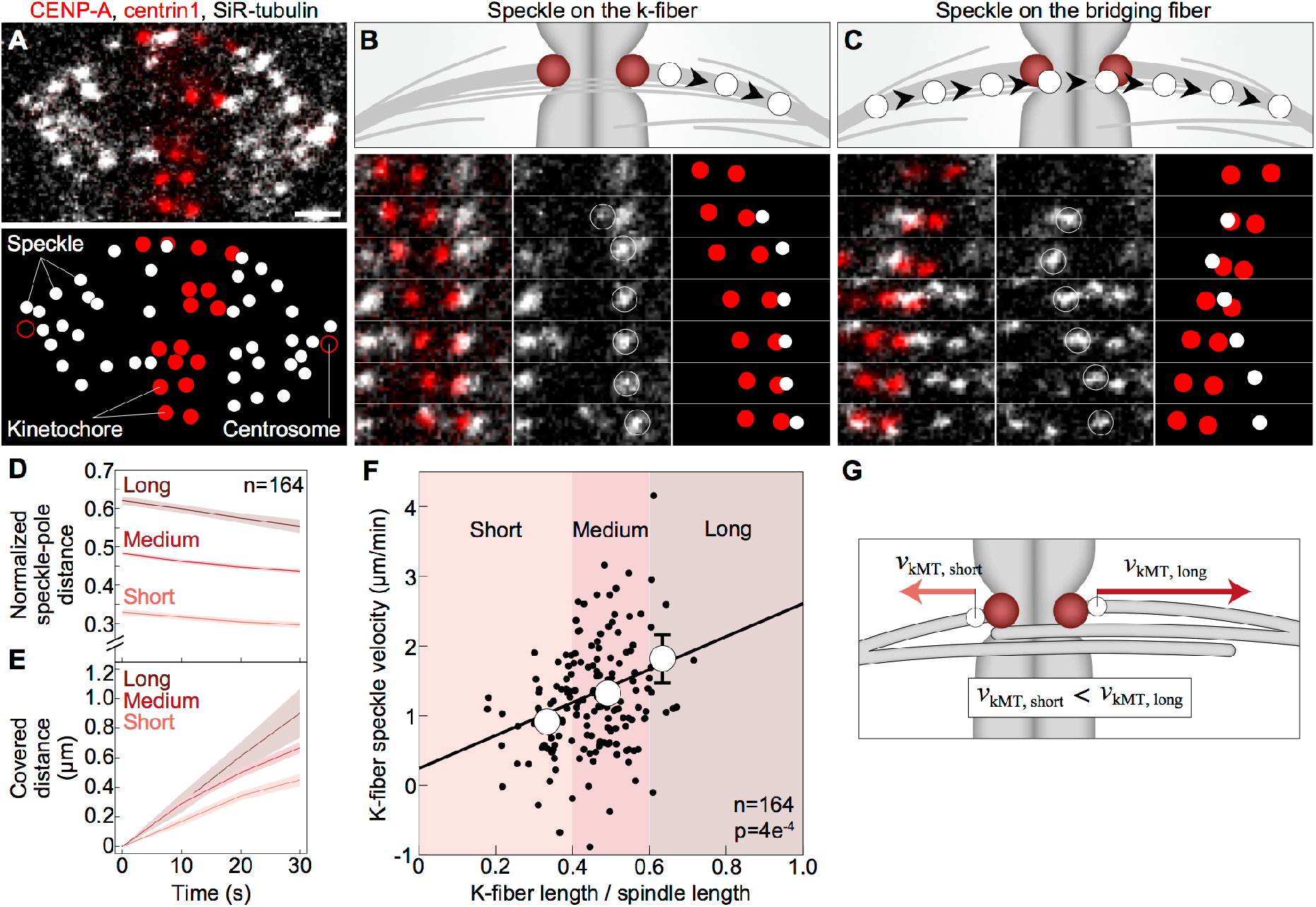
Poleward flux promotes kinetochore movement towards the spindle midplane. (**A-C**) Speckle microscopy assay for measurement of poleward flux of individual microtubules. (**A**) Spindle in a RPE1 cell stably expressing CENP-A-GFP and centrin1-GFP (red) stained with 1 nM SiR-tubulin dye which appears as distinct speckles marking individual microtubules (grey). Scale bar: 2 µm. (**B**) Scheme of a speckle originating at the kinetochore defined as the one marking a k-fiber microtubule (top). Montage over time demonstrating the movement of the speckle belonging to the k-fiber microtubule (bottom). Left shows merge, middle shows SiR-tubulin channel with encircled speckle, right shows schematic of kinetochores (red) and speckle (white) positions. (**C**) Scheme of a speckle passing the region between sister kinetochores, moving close to the kinetochores, defined as the one marking a microtubule within the bridging fiber (top). Montage over time demonstrating the movement of the speckle belonging to the bridging fiber microtubule. Legend as in B. (**D**) Speckle-pole distance over time divided by spindle length for k-fibers classified as short, medium, and long, according to the k-fiber length being smaller than 0.4, between 0.4 and 0.6, and larger than 0.6 of the spindle length, respectively. (**E**) Change in speckle-pole distance over time for speckles within groups as in D. (**F**) Poleward velocity of k-fiber speckles within groups as in D depending on its relative starting speckle-pole distance. (**G**) Scheme of speckles on longer and shorter k-fiber, where the speckle on the longer k-fiber fluxes faster than the speckle on the shorter k-fiber.

To identify the speckles that are localized on kinetochore or bridging microtubules, we follow the position of their first appearance and their subsequent movement. The speckles that originate close to a kinetochore, at the pole-facing side, were defined as those on a kinetochore microtubule (**Fig. 2 B**). The speckles that appear on one side of a pair of sister kinetochores, pass the region between them, and end up on the other side, were defined as those on a bridging microtubule (**Fig. 2 C**). All other speckles in the spindle region between the centrosomes, for which we cannot determine the type of microtubule they belong to, we refer to as "other" speckles (**Fig. S2 A, B**). We tracked individual speckles (**Fig. S2 C**) together with the spindle poles marked by centrioles, and calculated poleward flux as the change of the speckle-to-pole distance over the first 30 seconds of their movement (**Table 1**). This assay allowed us to study the movement of kinetochore and bridging microtubules with respect to the poles and to each other.

**Table 1.**
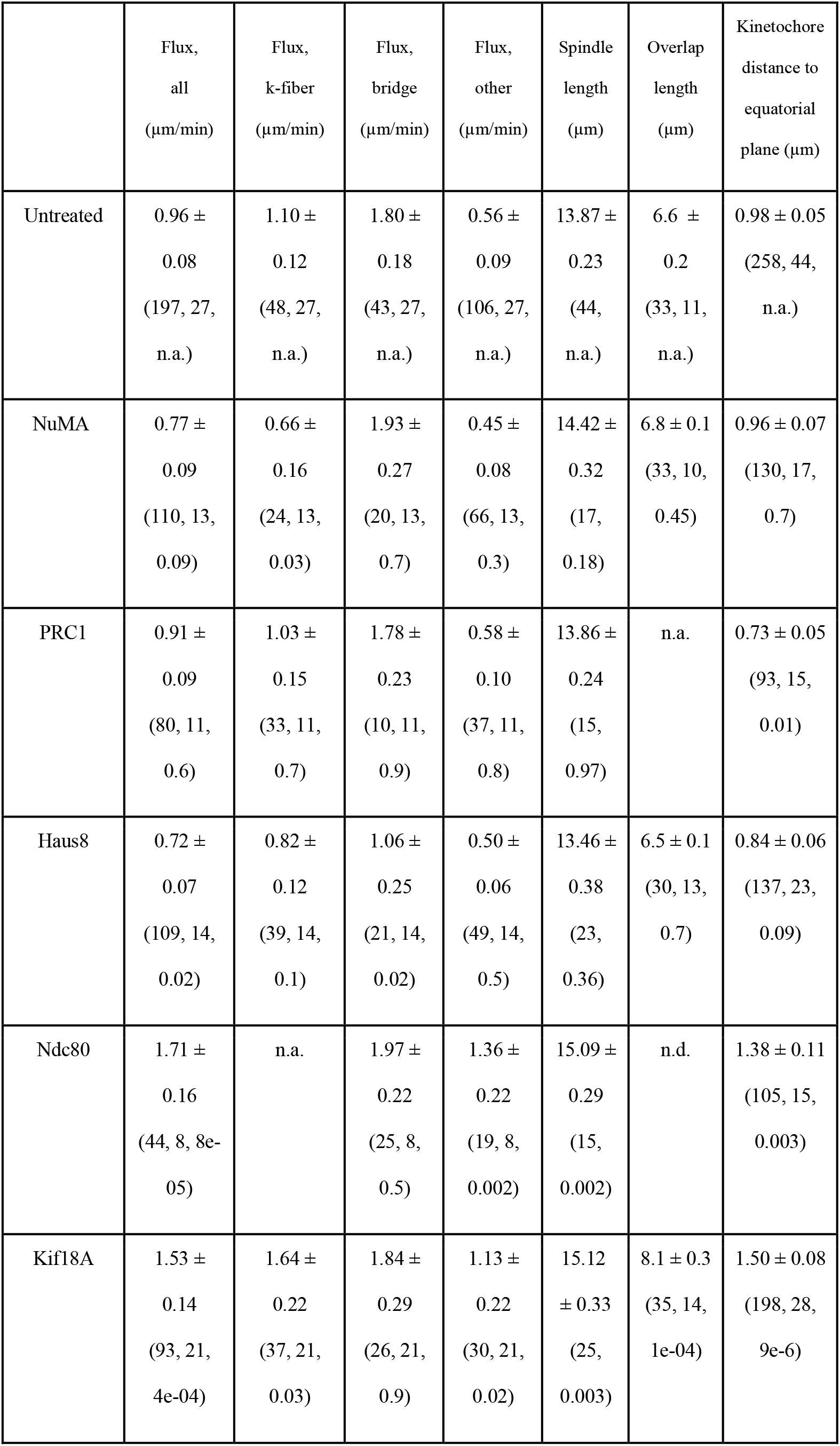
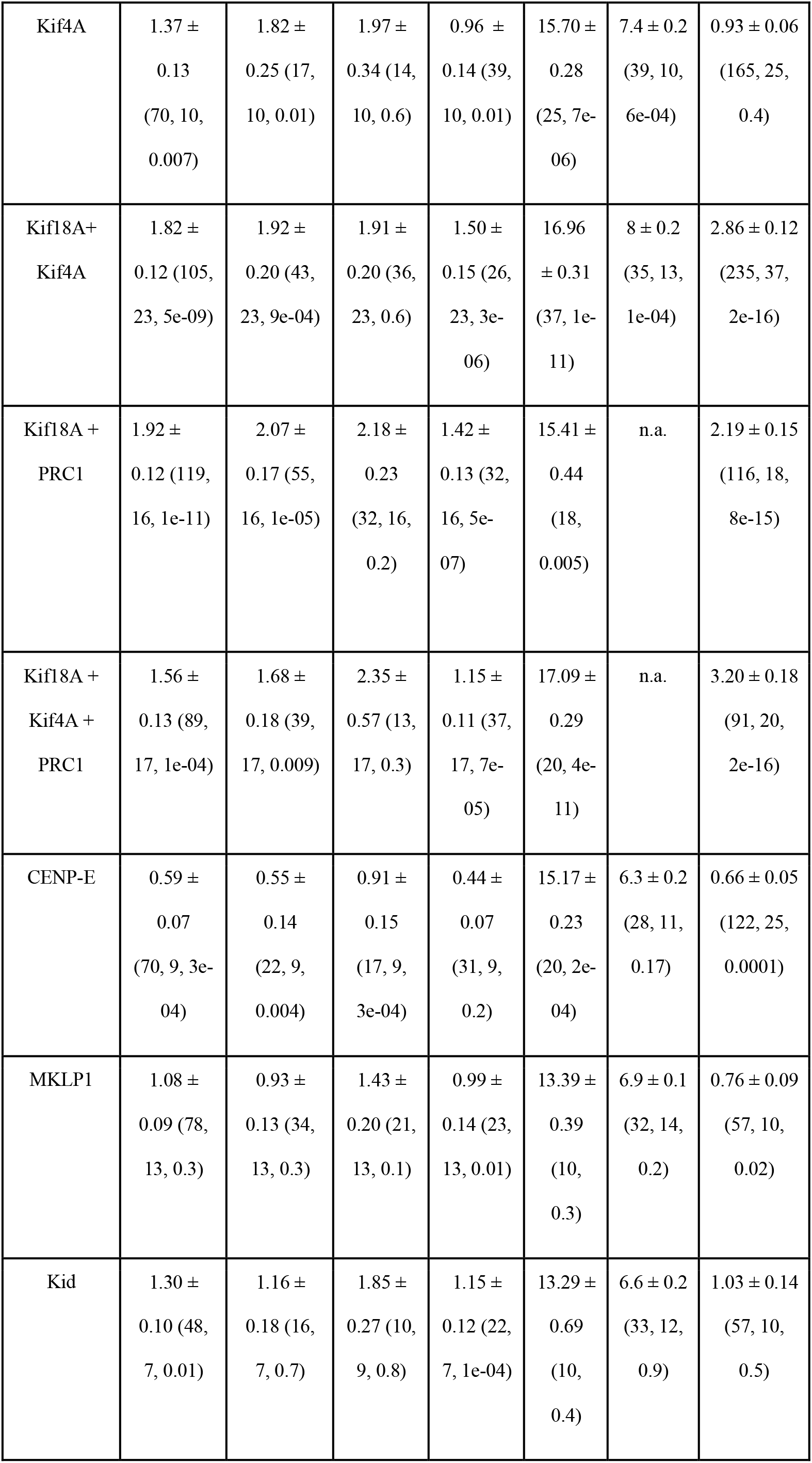
Measurements of flux, spindle and kinetochore parameters. Values are given as mean ± sem. The numbers in brackets denote the number of measurements (number of speckles for flux measurements or number of kinetochore pairs; for spindle length this number is not given because it is equal to the number of cells), number of cells, and p-value from a t-test for comparison with untreated cells. *n.a.*, not applicable; *n.d.*, not determined.

To explore the relevance of the model to kinetochore alignment, we used this assay in unperturbed cells and after a set of perturbations in which we depleted candidate microtubule-associated proteins by siRNA. We depleted motor proteins that are known to be involved in kinetochore alignment and/or localize to the bridging fiber: Kif18A, Kif4A, Kid, CENP-E, and MKLP1, and non-motor proteins that are important for k-fiber and bridging fiber integrity and their crosslinking: PRC1, NuMA, and Haus8 (Maiato et al., 2017; Tolić and Pavin, 2021) (see **Fig. S3 A-H** for depletion efficiency). For all these treatments, we analyzed the poleward flux of bridging and k-fibers (**Fig. S4 A-L**), kinetochore positions and the length of anti-parallel overlaps (**Table 1**). Although these treatments most likely also affect other aspects of the spindle architecture and dynamics, we expect to identify general interdependence between flux dynamics and kinetochore centering.

**Figure 3.**
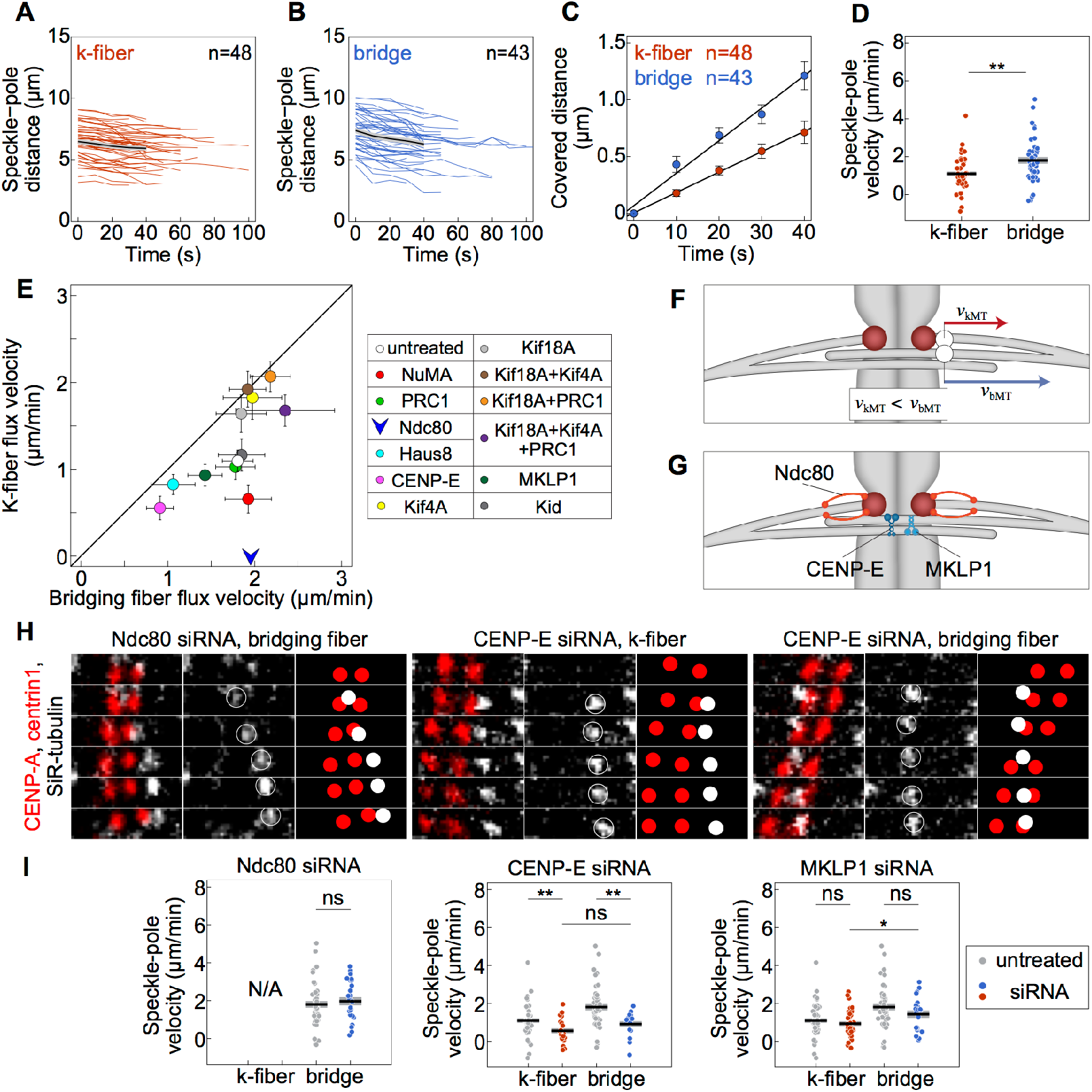
Bridging microtubules flux faster than kinetochore microtubules. Distance between k-fiber (**A**) and bridging fiber (**B**) speckles from the corresponding pole over time in untreated cells. Colored lines show individual speckles. Black line; mean. Grey area; s.e.m. (**C**) Change in speckle-pole distance over time for speckles within k-fibers and bridging fibers in untreated cells. (**D**) Poleward velocity of the k-fiber and bridging fiber speckles. Each dot corresponds to an individual speckle. Black line; mean. Grey area; s.e.m. (**E**) Poleward velocity of the k-fiber versus poleward velocity of the bridging fiber (left). siRNA treatments are color-coded as shown in the legend (right). Note that Ndc80-depleted cells are shown as arrow because poleward velocity of k-fibers could not be assessed. (**F**) Scheme showing that a speckle within the bridging fiber fluxes faster than a speckle within the k-fiber. (**G**) Scheme of localization of Ndc80, CENP-E, and MKLP1. (**H**) Montage over time demonstrating the movement of a speckle belonging to the bridging fiber in Ndc80 siRNA treatment (left) and k-fiber (middle) and bridging fiber (right) in CENP-E siRNA treatment. Legend as in **Fig. 1 B**. (**I**) Speckle-pole velocities after depletion of Ndc80, CENP-E, and MKLP1 (left to right). Graphs show poleward velocity of the speckles in siRNA-treated (red: k-fiber, blue: bridging fiber) and untreated (grey) cells. Black line; mean. Grey area; s.e.m. Statistical analysis, t-test. p-value legend: < 0.0001 (****), 0.0001 to 0.001 (***), 0.001 to 0.01 (**), 0.01 to 0.05 (*), ≥ 0.05 (ns).

**Figure 4.**
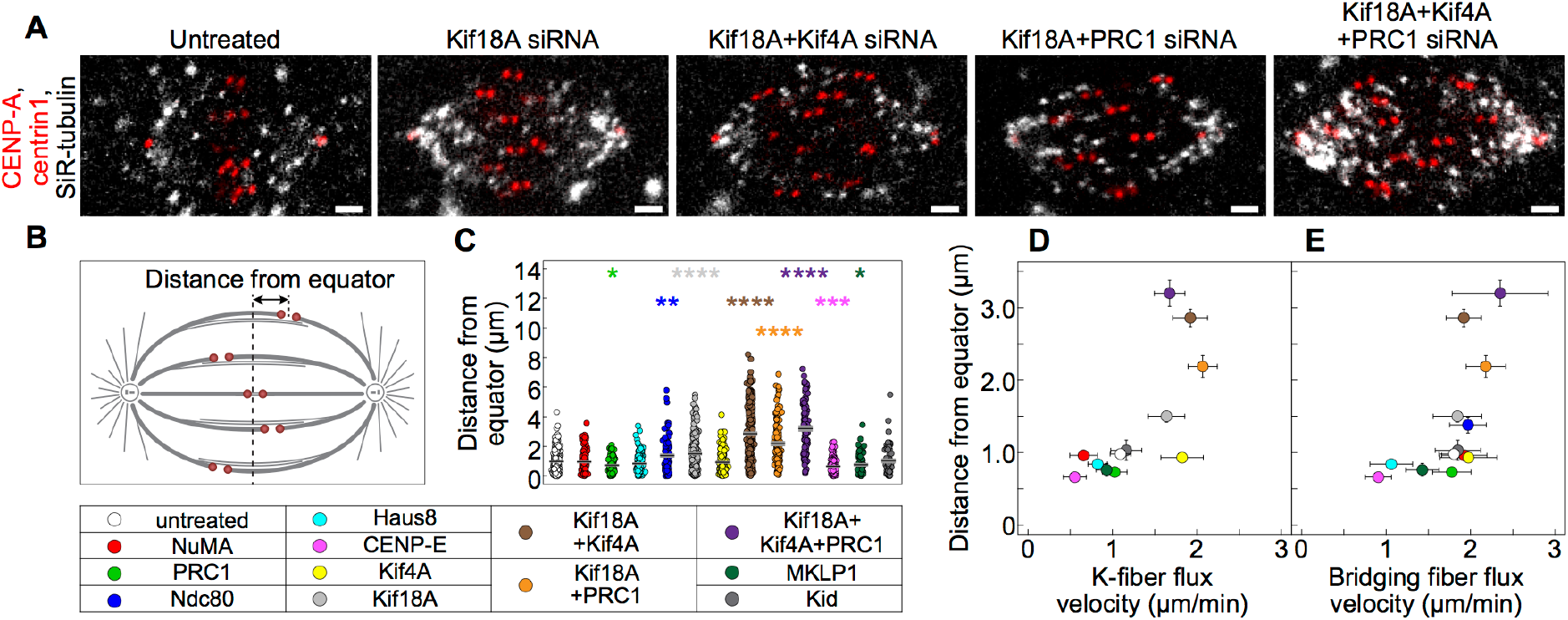
Kinetochore alignment depends on the flux velocity of k-fibers. (**A**) Spindle in a RPE1 cell stably expressing CENP-A-GFP and centrin1-GFP (red) stained with 1 nM SiR-tubulin dye. Cells are untreated, depleted for Kif18A, Kif18A and Kif4A, Kif18A and PRC1, and Kif18A, Kif4A and PRC1 (from left to right). Scale bar: 2 µm. (**B**) Scheme shows that the distance from equator was measured as the distance between sister kinetochore midpoint and the equatorial plane. (**C**) Kinetochore distance from equator in untreated and siRNA-treated cells. Each treatment is compared with untreated cells. Kinetochore distance from equator versus k-fiber (**D**) and bridging fiber (**E**) flux velocity in untreated and siRNA-treated cells. Treatments in (**C**)-(**E**) are color-coded according to the legend at the bottom. Statistical analysis, Mann-Whitney test; p-values as in **Fig. 3**.

### Longer kinetochore fiber undergoes flux at a higher velocity than the shorter one

The central prediction of the flux-driven centering model is that kinetochore centering relies on a difference in the flux velocity of sister k-fibers, where the flux of the longer k-fiber is faster than the flux of the shorter one. To explore whether this prediction holds in real spindles, we compared the flux of k-fibers of different lengths by using our speckle microscopy assay. The speckles on k-fibers, defined as those originating close to a kinetochore, were located at various distances from the pole, which correspond to the k-fiber length.

Strikingly, the k-fiber poleward velocity increased with an increasing k-fiber length in untreated cells (p = 4e-04, n = 164, **Fig. 2 D-F**). The same trend was observed when the k-fibers were divided into 3 groups: short, medium, and long, as those with a length smaller than 0.4, between 0.4 and 0.6, and larger than 0.6 of the spindle length, respectively. Short k-fibers had a flux of 0.91 ± 0.08 µm/min (n = 51 speckles from 68 cells), whereas the flux of long k-fibers was significantly faster, 1.81 ± 0.34 µm/min (n = 11 speckles from 68 cells, p = 3e-04), and the flux of medium k-fibers was between these values. The average poleward flux velocity of all speckles on k-fibers was 1.10 ± 0.12 µm/min (n = 48 speckles from 27 cells), which is similar to the flux rate previously measured by tubulin photoactivation on kinetochore fibers in RPE1 cells expressing photoactivatable-GFP-α-tubulin (Dudka et al., 2018), supporting our criteria for identification of speckles on k-fibers. Taken together, our experiments reveal that longer k-fibers flux faster than shorter ones, which is a key feature of the flux-driven centering mechanism (**Fig. 2 G**).

### Bridging microtubules undergo poleward flux at a higher velocity than kinetochore microtubules

In the flux-driven centering mechanism, motors within the bridging fiber drive the flux of bridging microtubules, and the interaction between the bridging and k-fibers generates the flux of k-fibers. However, the tension between sister kinetochores opposes the flux of k-fibers, making it slower than the flux of the bridging fiber. This difference between the bridging and k-fiber flux is found in the model for various parameters (**Fig. 1 D, F, H, J**), so we asked if the same feature is also observed in experiments.

Remarkably, speckles on the bridging microtubules moved poleward at a velocity of 1.80 ± 0.18 µm/min in untreated cells (n = 43 speckles from 27 cells), which is significantly faster than for the speckles on kinetochore microtubules (p = 0.001) (**Fig. 3 A-D; Table 1**). Because our experiments provide the first measurement of poleward flux of bridging microtubules in human spindles, we decided to validate our method of identification of speckles in the bridging fiber. We used PRC1 siRNA, which is known to specifically reduce the number of bridging microtubule to ∼50% of the original number (Jagrić et al., 2021; Polak et al., 2017). In cells treated with PRC1 siRNA, we observed 10 speckles on bridging microtubules in 11 cells, which is roughly a 2-fold reduction in comparison with untreated cells, where 43 speckles were observed in 27 cells, providing support for our method of identifications of speckles on bridging fibers (**Fig. S5 A**).

**Figure 5.**
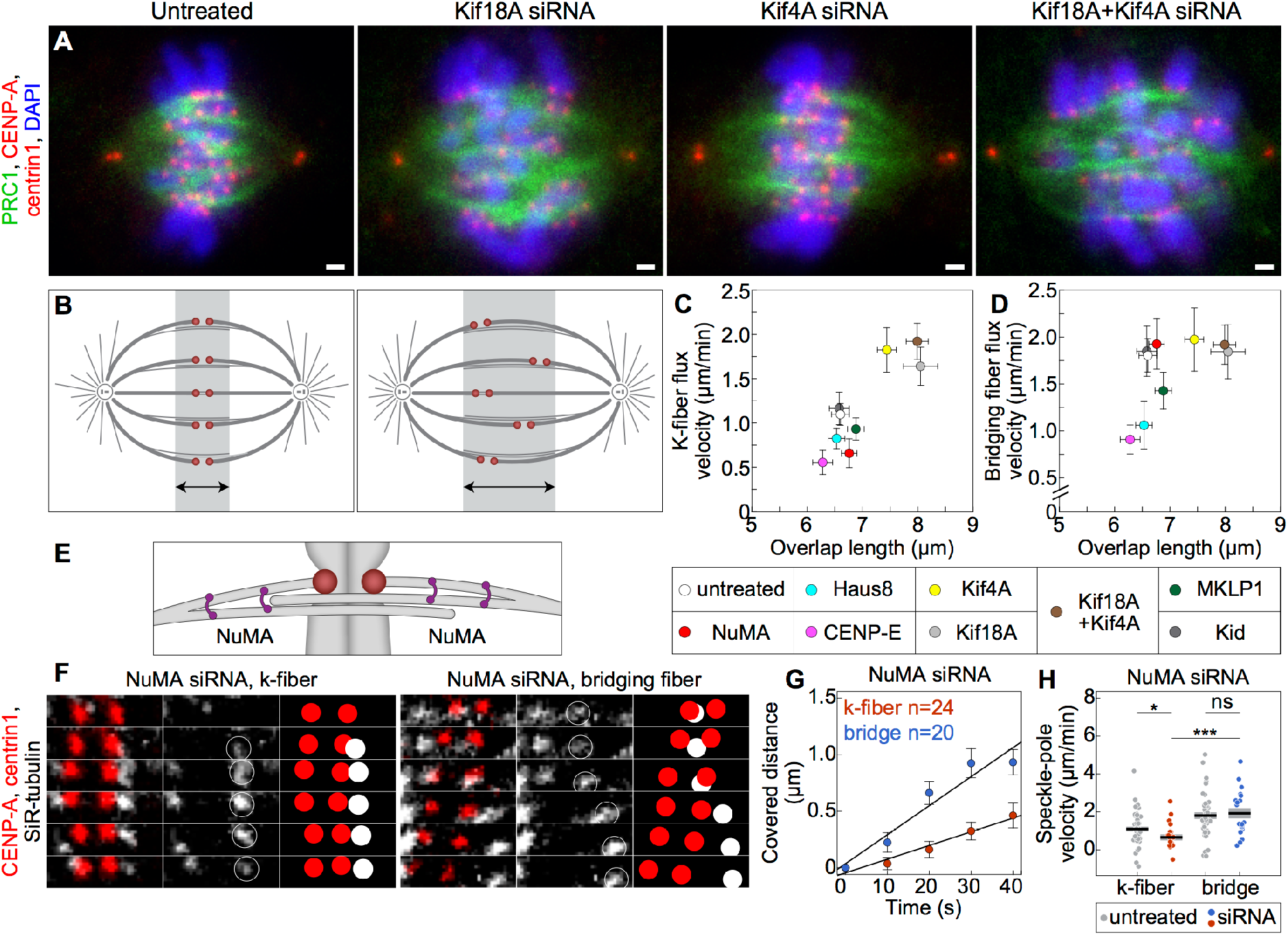
Coupling between bridging and k-fibers controls k-fiber flux velocity. (**A**) Fixed spindles in RPE1 cells stably expressing CENP-A-GFP and centrin1-GFP (red) in untreated, Kif18A siRNA, Kif4A siRNA, and Kif18A and Kif4A siRNA treated cells (from left to right), immunostained for endogenous PRC1 (AF-594, green) and stained with DAPI (blue). Images are sum intensity projections of five z-planes. Scale bar: 1 µm. (**B**) Scheme shows that spindles with shorter (left) and longer (right) overlap regions have better (left) and worse (right) kinetochore alignment at the spindle equator, respectively. K-fiber (**C**) and bridging fiber (**D**) flux velocity versus PRC1-labeled overlap length. Treatments are color-coded as shown in the legend below. (**E**) Scheme of NuMA localization. (**F**) Montage over time demonstrating the movement of a speckle belonging to the k-fiber (left) and bridging fiber (right) in NuMA siRNA treatment. Legend as in **Fig. 1 B**. (**G**) Change in speckle-pole distance over time for speckles within bridging and k-fibers in cells treated with NuMA siRNA. (**H**) Poleward velocity of the speckles in NuMA siRNA-treated (red: k-fiber, blue: bridging fiber) and untreated (grey) cells. Black line; mean. Grey area; s.e.m. Statistical analysis, t-test. p-value as in **Fig. 3**.

The observed rate of bridging microtubule poleward flux implies that the antiparallel bridging microtubules slide apart with respect to each other at twice the rate of their poleward flux, i.e. 3.6 ± 0.3 µm/min, given that the spindle length is constant during metaphase. This rate is comparable to the sliding rate of bridging microtubules in early anaphase measured by tubulin photoactivation, which is roughly 4.5 µm/min (Vukušić et al., 2021), suggesting that the bridging microtubule sliding may be driven by a similar mechanism in metaphase and early anaphase.

To explore the relationship between the bridging and k-fiber flux under various perturbations of the spindle, we measured the flux after a set of depletions of spindle proteins given in **Table 1**. Strikingly, the flux of bridging fibers was faster than or equal to the flux of k-fibers across the treatments (**Fig. 3 E**). Thus, the relationship between the bridging and k-fiber flux predicted by the model holds also in altered spindles with changed flux velocities (**Fig. 3 F**). Because a difference in flux velocities between k-fibers and bridging fibers is the signature of the flux-driven centering mechanism, our experimental findings in untreated cells as well as after various treatments suggest that this mechanism is relevant for kinetochore centering.

To explore to what extent k-fibers affect the sliding of bridging microtubules, we depleted Ndc80, the main coupler of kinetochores to microtubule ends (**Fig. 3 G; Movie 3**) (Cheeseman et al., 2006; Cheeseman and Desai, 2008). As expected, we did not detect speckles on k-fibers, i.e., those at the pole-facing side of the kinetochore, after Ndc80 depletion (n=8 cells). We found that the speckles on bridging microtubules fluxed at a similar velocity as in untreated cells (**Fig. 3 H, I; Table 1**), suggesting that sliding of bridging microtubules is largely unaffected by k-fibers and that the poleward flux is generated within the bridging fiber. In contrast to Ndc80, depletion of CENP-E or MKLP1 led to a decrease in bridging flux velocity of ∼50% and ∼20%, respectively (**Fig. 3 H, I; Movie 3; Table 1**), suggesting that these motors contribute to antiparallel sliding within bridging fiber overlaps (**Fig. 3 G)**.

### Kinetochore centering efficiency depends on the flux velocity of k-fibers

At the core of this centering mechanism is that the shorter k-fiber has slower flux than the longer sister k-fiber, generating a flux difference that moves the off-centered kinetochores towards the spindle center. This difference in flux requires the average k-fiber flux to be slower than the bridging fiber flux. Thus, centering is more efficient when k-fiber flux is slower than the bridging fiber flux, allowing for sliding of k-fibers along bridging fibers. This relationship between the centering efficiency and the k-fiber flux velocity is an important testable prediction of the model (**Fig. 1 E-J**).

To compare our experiments with the model, we quantified kinetochore centering efficiency by measuring the distances of sister kinetochore midpoints from the equatorial plane of the spindle (**Fig. 4 A-C; Movie 4**) and explored how this distance depends on the k-fiber flux velocity across all treatments (**Fig. 4 D**). To interpret this dependence, we focus on the general trend in these data, which shows that faster k-fiber flux is associated with worse centering. The treatments with the k-fiber flux velocity similar to that of untreated cells show efficient centering comparable to untreated cells. In contrast, treatments with faster k-fiber flux show worse centering. Among these treatments, the centering efficiency was highly variable even though the k-fiber flux velocities were similar, which is likely due to other mechanisms that operate in addition to those related to the flux (see **Discussion**). Unlike k-fiber flux, the bridging fiber flux remained unchanged in the treatments with worse centering (**Fig. 4 E**).

As a control for proper attachment of misaligned kinetochores we imaged astrin, which binds to end-on attached kinetochores (Shrestha et al., 2017), and found it localized at all kinetochores including those that were highly off-centered (**Fig. S5 B**). This suggests that the reason for off-centering was not lack of kinetochore biorientation. We also note that the observed worse centering after combined depletion of Kif18A and Kif4A in comparison with Kif18A depletion differs from a previous report (Stumpff et al., 2012). This difference is not due to the use of different cell lines as we obtained similar results on HeLa and U2OS cells as on RPE1 (**Fig. S5 C, D**), but likely related to a different effect of the double depletion on spindle length. Taken together, our experiments and the model suggest that kinetochores are better centered when the k-fiber flux is markedly slower than the bridging fiber flux, allowing for sliding of k-fibers along bridging fibers and thus the movement of the center of sister k-fibers towards the spindle center.

### Longer overlaps of antiparallel microtubules lead to an increase in the k-fiber flux velocity to the bridging fiber flux velocity

Our experiments have shown that an increased flux velocity of k-fibers is related to less efficient kinetochore centering. What caused this speeding up of the k-fiber flux in the treatments with misaligned kinetochores? The model suggests that changes in the overlap length can lead to changes in flux velocities (**Fig. 1 F**), even though these two quantities are not obviously correlated.

To explore this intriguing relationship, we measured the overlap length by measuring the length of PRC1-labeled regions in all the treatments except those where PRC1 was depleted (**Fig. 5 A, B; Fig. S5 E**). Out of these treatments, overlaps were longer after depletion of Kif18A or Kif4A, in agreement with previous results (Jagrić et al., 2021), and after a combined depletion of Kif18A and Kif4A (**Table 1;** see **Fig. S5 F** for HeLa and U2OS cells). These treatments specifically increased the flux velocity of k-fibers without changing the flux of bridging fibers, resulting in k-fibers fluxing at ∼90% of the bridging fiber flux velocity (**Fig. 5 C, D; Table 1**). For comparison, in untreated cells k-fibers flux at ∼60% of the bridging fiber flux velocity (**Table 1**). Thus, these experiments reveal a relationship between the overlap length and the k-fiber flux velocity, suggesting that the sliding forces generated within the bridging fiber are transferred to the k-fibers through the antiparallel overlaps between these two types of fibers.

### The difference in the flux of bridging and k-fibers increases for smaller concentrations of passive crosslinkers

The sliding forces from the bridging fiber are transmitted to the k-fibers not only through the antiparallel overlaps, but also through the regions of parallel overlaps, where the bridging and kinetochore microtubules extending from the same spindle half are linked together by passive crosslinkers (see **Fig. 1A**). Thus, reducing the amount of passive crosslinkers should result in reduced force transmitted from the bridging to the k-fibers and consequently in slower flux of k-fibers, as predicted by the model (**Fig. 1 H**).

To explore the role of passive crosslinkers in the parallel overlaps of bridging and kinetochore microtubules, we chose NuMA as a candidate because it is required for local load-bearing in the spindle (**Fig. 5 E**) (Elting et al., 2017) and for synchronous microtubule flux across the spindle (Steblyanko et al., 2020). After depletion of NuMA by siRNA (**Fig. 5 F; Fig. S3 G; Movie 5**), we found that the flux velocity of kinetochore microtubules decreased by ∼40% (from 1.10 ± 0.12 in untreated cells to 0.66 ± 0.16 µm/min after NuMA depletion, **Fig. 5 G, H**; **Fig. S4 I; Table 1**). On the contrary, the flux velocity of bridging fibers did not change significantly (**Fig. 5 G, H**), thus the difference compared to the k-fiber velocity increased. Because the model predicts a larger difference in flux velocities for fewer passive crosslinkers (**Fig. 1 H**), these results support the idea that NuMA acts as a passive crosslinker transmitting the sliding forces from the bridging fiber onto the associated k-fibers through their parallel overlaps.

## DISCUSSION

### Flux-driven centering model explains kinetochore alignment at the equatorial plane

Based on our model and the results from speckle microscopy that allowed us to measure the relative movements of kinetochore and bridging microtubules, we propose that microtubule poleward flux promotes kinetochore centering. Motor proteins within the overlaps of k-fibers and bridging microtubules generate sliding forces proportional to the overlap length, which drive poleward flux of the k-fiber. Thus, the flux of a longer sister k-fiber is faster than the flux of the shorter one, resulting in a tug-of-war in which the longer k-fiber wins and the kinetochores move towards the spindle center (**Fig. 6 A**). This key feature of the centering mechanism was indeed observed in our experiments where we tracked speckles on k-fibers of different length.

**Figure 6.**
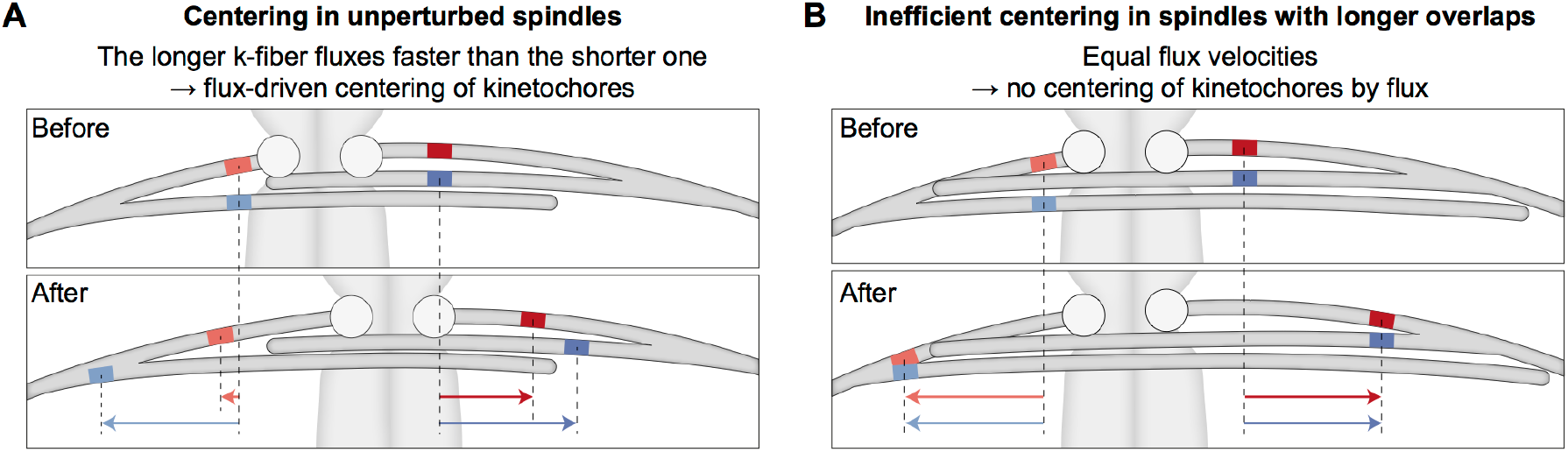
Mechanism by which poleward flux promotes kinetochore centering. (**A**) A pair of kinetochores (circles) is displaced towards the left (top). To visualize relative movements of the microtubules, four marks are shown (red and blue). Over time (bottom), the marks on the bridging microtubules move poleward by a similar distance (arrows), whereas the marks on the k-fibers move more slowly due to imperfect coupling between the bridging and k-fibers. Importantly, the longer k-fiber on the right side has a longer overlap with the bridging fiber and thus the coupling is stronger, leading to a higher flux velocity of this fiber in comparison with the shorter k-fiber, which in turn results in the movement of the kinetochores towards the spindle center. (**B**) If the coupling between the k-fibers and the bridging fiber is too strong, such as in cases when the antiparallel overlaps are excessively long, the k-fibers flux velocity becomes similar to the velocity of the bridging fiber. Thus, k-fibers do not slide with respect to the bridging fiber, resulting in chromosome misalignment.

The flux-driven centering mechanism proposed here and the previously introduced centering forces based on length-dependent suppression of k-fiber dynamics (Gardner et al., 2008; Mary et al., 2015; Gergely et al., 2016; Klemm et al., 2018) as well as on polar ejection forces (Joglekar and Hunt, 2002; Civelekoglu-Scholey et al., 2006, 2013; Armond et al., 2015) are conceptually independent. Yet, proteins such as Kif18A and Kif4A may be involved in more than one mechanism. These centering mechanisms may work together but with different efficiency depending on the cell type and the stage of spindle assembly. Due to the complexity of the spindle, it is hard to identify the contribution of each mechanism by using only experimental approaches, but future theoretical studies combining different centering mechanisms may help to identify the dominant mechanism.

### Kinetochore fiber flux is driven by interactions with the bridging fiber

By developing a speckle microscopy assay based on low doses of SiR-tubulin to distinguish kinetochore and bridging microtubules, our work demonstrated that bridging microtubules undergo poleward flux and this flux is faster than that of kinetochore microtubules. In contrast to metaphase, k-fibers and bridging fibers slide together at a similar rate in early anaphase (Vukušić et al., 2017). This difference is most likely due the tension between sister kinetochores, which is present in metaphase when sister kinetochores are linked by chromatin, but not in anaphase when this link vanishes. By using various siRNA-mediated protein depletions, we were able to increase or decrease the difference in the metaphase flux velocities, but the k-fiber flux remained slower or equal to the bridging fiber flux, suggesting that motor proteins and crosslinkers regulate the relationship between these two velocities. Interestingly, slower flux of kinetochore microtubules than adjacent non-kinetochore ones was observed in *Xenopus* egg extracts (Maddox et al., 2003; Yang et al., 2008) and crane-fly spermatocytes (LaFountain et al., 2004), indicating that the relationship between the flux of these two sets of microtubules is conserved across organisms whose spindles undergo flux. As the flux-driven centering mechanism relies on this difference, we propose that flux could promote chromosome centering in a variety of organisms.

Two flux velocities have been observed also in human U2OS cells, where the subset of microtubules fluxing faster than kinetochore microtubules was associated with the γ-tubulin ring complex (γTuRC) (Lecland and Lüders, 2014). γTuRC is recruited to microtubules by the augmin complex to nucleate new microtubules (Kamasaki et al., 2013; Uehara et al., 2009; David et al., 2019; Goshima et al., 2008), including nucleation of bridging microtubules along k-fibers (Manenica et al., 2020; O’Toole et al., 2020). Thus, the fast flux of bridging microtubules in comparison with k-fibers measured here likely corresponds to the fast γTuRC- nucleated microtubule fraction. Our observation that the flux of bridging microtubules slowed down after depletion of the augmin subunit Haus8 supports this conclusion.

In spindles without k-fibers, i.e, after Ndc80 depletion, the overall flux was faster than in untreated cells, in agreement with a recent study (Steblyanko et al., 2020). This velocity was similar to the flux velocity of bridging fibers in untreated cells, showing that bridging fiber flux is largely unaffected by k-fibers. This result, together with the faster flux of bridging fibers in comparison with k-fibers across treatments, suggests that bridging fibers drive poleward flux of k-fibers. The forces driving poleward flux have been debated, where the dominant forces are thought to be either at the spindle pole (Rogers et al., 2004; Ganem et al., 2005) or within interpolar microtubules (Miyamoto et al., 2004; Brust-Mascher et al., 2009; Matos et al., 2009). Thus, our experiments are in agreement with the latter possibility and support the assumption of our model that the leading forces are generated within antiparallel overlaps.

We found that the depletion of NuMA, a passive crosslinker of microtubules (Elting et al., 2017), decreases poleward flux of kinetochore microtubules without affecting the flux of bridging microtubules. As our model predicts slower k-fiber flux for decreased amount of passive crosslinkers, our experiments together with the model suggests that NuMA transmits the force from the bridging onto the k-fibers. Interestingly, NuMA depletion was shown to cause asynchrony of microtubule poleward flux (Steblyanko et al., 2020), implying that NuMA crosslinks neighboring k-fibers and synchronizes their flux. Because our flux-driven centering mechanism relies on k-fiber flux velocities, we suggest that the correlated movement of neighboring kinetochore pairs (Vladimirou et al., 2013) reflects the synchrony in poleward flux of neighboring k-fibers. In addition to NuMA, bridging microtubules may promote synchrony in k-fiber flux as bridging microtubules were shown to fan out at their ends and interact with neighboring k-fibers (O’Toole et al., 2020).

Depletion of PRC1 did not change the flux velocity of bridging or kinetochore microtubules, in agreement with a previous study (Steblyanko et al., 2020). As PRC1 depletion leads to ∼50% decrease of the number of microtubules in the bridging fiber (Jagrić et al., 2021), our result suggests that the remaining bridging microtubules are sufficient to generate flux. In contrast to PRC1, CENP-E and MKLP1 depletion led to a decrease in bridging flux velocity. Given that CENP-E localizes to the bridging fibers in metaphase (Steblyanko et al., 2020; Jagrić et al., 2021), together with MKLP1 (Jagrić et al., 2021), these motors may contribute to antiparallel sliding within bridging fiber overlaps. The effect of CENP-E depletion may also be due to the role of CENP-E in targeting CLASPs, which promote flux, to kinetochores (Maiato et al., 2005; Maffini et al., 2009; Girão et al., 2020). Even though in our model the sliding of antiparallel microtubules is driven by motor proteins with the same physical properties, previous work (Steblyanko et al., 2020) and our experiments suggest that multiple different motor proteins contribute to sliding in real spindles. These motors may have different physical properties and thus bring new phenomena to the flux-driven centering mechanism by affecting sliding forces.

### Chromosome alignment depends on the overlap length of bridging microtubules

In our model, the length of antiparallel overlaps determines the flux velocity of k-fibers and kinetochore centering. If overlaps are longer, the coupling between bridging and k-fibers increases, leading to an increase in the k-fiber flux velocity to the bridging fiber flux velocity. This prevents sliding of k-fibers along the bridging fiber and results in worse kinetochore centering (**Fig. 6 B**).

Our experiments with depletions of spindle proteins revealed a trend where kinetochores are misaligned when overlaps are longer and k-fiber flux is faster. For example, depletion of Kif18A resulted in extended overlaps, faster k-fiber flux, and misaligned chromosomes. We propose that Kif18A regulates chromosome alignment by the flux-driven centering mechanism, where this motor protein controls k-fiber flux through the regulation of the length of antiparallel overlaps between bridging and k-fibers. Previous work showing that Kif18A is localized within bridging fibers in addition to k-fibers (Jagrić et al., 2021) supports this possibility. This mechanism may work together with the known mechanism where Kif18A suppresses the dynamics of longer k-fibers at their plus ends (Stumpff et al., 2012; Du et al., 2010). To understand the roles of the Kif18A motors localized within bridging fibers versus those on k-fibers in chromosome alignment, it will be crucial to develop approaches based on separation-of-function experiments.

In contrast to depletions of Kif18A alone or in combination with Kif4A and PRC1, depletion of Kif4A alone showed no effect on chromosome alignment even though the overlaps of bridging microtubules were extended and the k-fiber flux was faster. This is possibly due to the activity of Kif18A at the k-fiber tips.

Whereas in our model the relationship between k-fiber flux and kinetochore centering was observed by varying a single parameter, experimental perturbations relied on depletion of motor proteins, which have multiple functions within the spindle. Thus, the interpretation in light of the model is not always straightforward. For example, a closer comparison between experiments and the model could be achieved by extending the model to include regulation of length of bridging and kinetochore microtubules by Kif18A, as in this case one could explore the resulting overlap length and kinetochore positions after changing Kif18A concentration in the model.

### Outlook

Our work revealed a new mechanism of kinetochore alignment where lateral length-dependent sliding forces that the bridging fiber exerts onto k-fibers promote the movement of kinetochores towards the spindle center. We have shown that this mechanism drives kinetochore alignment during metaphase, but the same mechanism may also be a crucial part of spindle assembly in prometaphase to promote chromosome congression to the metaphase plate. Given that spindles in prometaphase undergo poleward flux (Steblyanko et al., 2020), future experiments will reveal whether and how it contributes to chromosome congression.

The physiological importance of chromosome alignment is in preventing lagging chromosomes and appearance of micronuclei, thereby promoting proper nuclear reformation and karyotype stability (Fonseca et al., 2019; Maiato et al., 2017). It will be interesting to explore the robustness of the flux-driven chromosome alignment and the resulting segregation fidelity in healthy cells. Even more importantly, future work should reveal what aberrations in this mechanism lead to errors in chromosome segregation in cells with unstable karyotypes in which misaligned chromosomes appear.

## METHODS

### Theory for kinetochore centering

The model describes a system consisting of two sister kinetochores, two microtubules representing the left and right k-fiber, which extend from the spindle edges to the kinetochores, and two bridging microtubules which extend from the edges and interdigitate in the middle (**Fig. 1 A, S1 E**). The positions of the sister kinetochores are denoted by 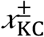 while the positions of k-fibers and bridging fibers are taken as arbitrary positions along their lattice and are denoted by 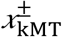 and 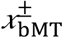 respectively. All these positions change in time *t* and their velocities are calculated as 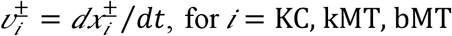 Hereon the superscripts *+* and − denote the right and left sides of the model, respectively. The length of the microtubule overlap within the bridging fiber is denoted *D*_0_ and the spindle length is denoted **L**_0_.

In order to calculate the movement of the kinetochores we first describe forces at them. The elastic connection between sister kinetochores is described by a force exerted by Hookean spring, 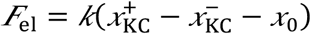 where *k* denotes the elastic coefficient and *x*_0_ the spring rest length. Kinetochores also interact with microtubules and the force exerted by microtubule plus end is described by 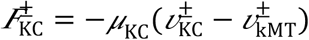 Here, *μ*_KC_ denotes the effective friction coefficient at the kinetochore. This description is a simplification of the force-velocity relationship measured for kinetochores (Akiyoshi et al., 2010). Because these forces, exerted by the elastic connection and the microtubule, are the only forces acting at kinetochores in our model, they balance each other,

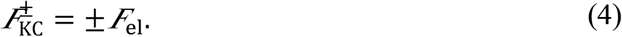

Equations (4) also include the balance of forces between sister kinetochores, 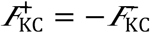.

The movement of the k-fiber is driven by forces exerted by molecular motors distributed along the k-fiber and bridging microtubule overlap, 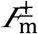 These forces are opposed by the damping force of the cross-linking proteins, 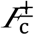 and by the force at the kinetochore,

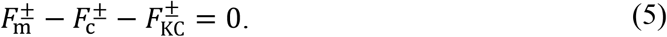

This expression is an application of equation (3) to left and right k-fibers.

The forces of the motors distributed along the antiparallel overlap of a k-fiber and a bridging fiber are described by equation (1), which for left and right sides reads 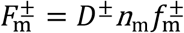 Force exerted by a single motor depends on the relative velocity of the k-fiber and the bridging fiber of opposite orientation and is described through a linear force-velocity relation 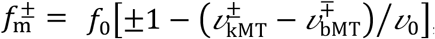 where **f**_0_ denotes the stall force and *v*_0_ the velocity without a load. The linear density of the motors is denoted *n*_m_, and the length of the antiparallel overlap of the bridging and k-fiber is given by 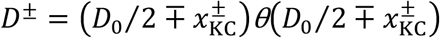 where *θ* is the Heaviside step function which ensures that the antiparallel overlap exists. The number of motors is given as 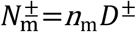.

The damping force of the crosslinking proteins is given by equation (2), which for left and right sides reads 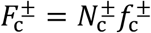, where the damping force of a single crosslinker 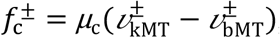 depends on the friction coefficient of a crosslinking protein, *μ*_c_, and the relative velocity of the k-fiber and the bridging fiber. The number of passive crosslinkers distributed along the parallel overlap of a k-fiber and a bridging fiber,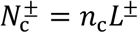 is calculated from linear density *n*_c_ and the length of the k-fiber 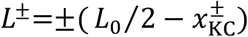.

The movement of the bridging microtubules is driven by the force exerted by motors distributed along the antiparallel overlap of bridging microtubules, **F**_bMT_. This force is opposed by the motor forces exerted along the antiparallel overlap of the bridging fiber and the k-fiber and by the damping force of the crosslinking proteins exerted along the parallel overlap of the bridging fiber and the k-fiber,

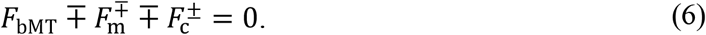

The force exerted in the overlap of bridging microtubules depends on their relative velocities, 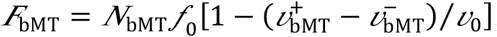 and the number of motors in the overlap of bridging microtubules, which is given as *N*_bMT_ = *n*_m_*D*_0_.

### Solution of the model

#### Approximations in the model

Even though the model can be solved as given, we introduce two approximations that apply, to a large extent, to the studied spindles. First, we neglect the difference in kinetochore velocities, 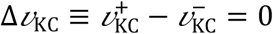 based on the following arguments. Interkinetochore velocity on a time scale relevant for centering of kinetochores 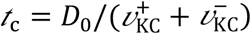 has an approximate value 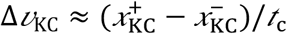. By applying (equation 4) on the left and right sides, the normalized interkinetochore velocity reads

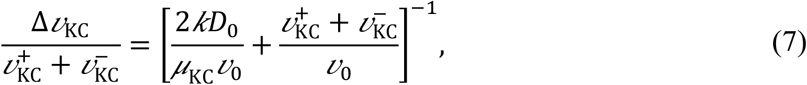

where we use 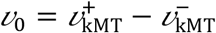 as an upper limit for k-fiber velocity difference. In the case of parameters that are relevant for our system, spring constant representing elasticity of the chromosomes, *k* = 100 pN/µm (Joglekar and Hunt, 2002), and *D*_0_, μ_KC_, and *v*_0_ from Figure 1B, the first term of the right-hand side of (equation 7) obeys 2*kD*_0_/μ_KC_*v*_0_ ≫ 1. In this limit the right-hand side of equation (7) approaches zero and the interkinetochore velocity can be neglected.

Second, we set the velocities of the bridging microtubules to the value 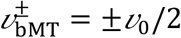 based on the following approximation. For kinetochores in the central position, the overlap regions on the left and right sides have the same length, *D*^+^ = *D*^−^ and *L*^+^= *L*^−^. By applying this symmetry to equations (6) we derive an expression:

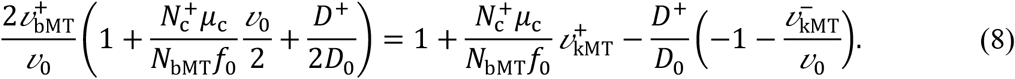

In the case where the contribution of the motors dominates over that of cross-linkers, 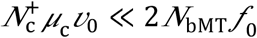, the second term in the bracket on the left side of the equation is much smaller than 1. Analogously, the second term on the right side can be neglected. Because the length of the antiparallel overlap between bridging and k-fibers is much smaller than the length of the overlap between bridging fibers, **D**^+^ ≪ **D**_0_, the third terms on both sides of the equation can be neglected. In this limit, Eq. (8) reduces to 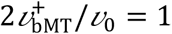.

#### Velocities of k-fibers and kinetochores

By applying these approximations, force-velocity relationship for individual motors, friction forces exerted by passive crosslinkers and kinetochores to equation (5) we derive expressions for k-fiber velocities:

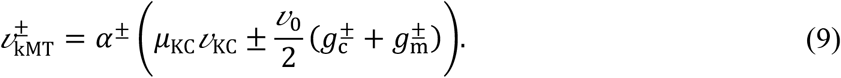

Here, to have shorter notations in our model, we define three symbols: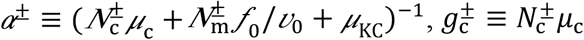, and 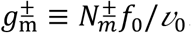.

Next, by combining the force balance on sister kinetochores, obtained from equations (4), with equations (5), we obtain a balance of forces between left and right k-fibers, which are exerted by motors and crosslinkers, 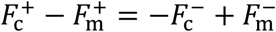. By applying force-velocity relationship for individual motors and friction forces exerted by passive crosslinkers to this expression, in combination with equation (9), we obtain a final expression for kinetochore velocities:

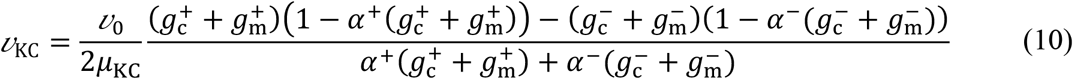

This final expression depends explicitly only on the geometry of the system. Thus, the positions of kinetochores can be calculated by integrating equation (10) over time.

### Choice of parameters

The value for the stall force of sliding motors *f*_0_ was taken from literature (Valentine et al., 2006) and the value for the sliding motor velocity without a load v_0_ was set to reproduce the measured velocity of the bridging microtubule and has a similar value to the microtubule sliding velocity that is driven by Eg5 motor proteins from *Xenopus laevis* (Hentrich and Surrey, 2010). The value for the effective friction coefficient μ_kc,_ is estimated as a ratio of the stall force at the kinetochore, which is 3 pN (Akiyoshi et al., 2010) and the polymerization velocity measured here, 0.1 µm/min. Two independent parameters, sliding motor density *n*_m_ and passive cross-linker friction multiplied by its density *n*_c_μ_c_, were considered as variable parameters and were varied over two orders of magnitude in order to explore the parameter space. Values for geometrical parameters, k-fiber length *L*_0_ and overlap length *D*_0_, were measured here.

## Experimental Methods

### Cell culture

hTERT-RPE-1 cell line with a stable expression of CENP-A-GFP and centrin1-GFP was a gift from Alexey Khodjakov (Wadsworth Center, New York State Department of Health, Albany, NY, USA). U2OS cell line expressing CENP-A-GFP was a gift from Marin Barisic (Danish Cancer Society Research Center, Copenhagen, Denmark) and Helder Maiato (Institute for Molecular Cell Biology, University of Porto, Portugal). HeLa cell line with a stable expression of EGFP-CENP-A was a gift from Andrew McAinsh (University of Warwick, Coventry, UK). Cells were maintained in Dulbecco’s Modified Eagle’s Medium (containing 1 g/L d-glucose, l-glutamine, pyruvate; Lonza) supplemented with 10% Fetal Bovine Serum (Sigma-Aldrich), 100 IU/mL penicilin (Lonza) and 100 mg/mL streptomycin (Lonza). Cells were grown at 37°C in a Galaxy 170s humidified incubator (Eppendorf) with a 5% CO2 atmosphere.

### RNA interference and transfection

One day before transfection, 120 000 cells were seeded on 35-mm glass coverslip dishes with 0.17-mm glass thickness (MatTek Corporation). siRNA constructs were diluted in Opti-MEM medium (Life Technologies) and transfection was performed with Lipofectamine RNAiMAX Reagent (Life Technologies) by following manufacturer’s protocol. Constructs and their final concentrations used were: 100 nM NuMA siRNA (sc-43978; Santa Cruz Biotechnology), 300 nM PRC1 siRNA (L-019491-00-0010; Dharmacon), 100 nM Ndc80 siRNA (HA12977117-004; Merck), 20 nM Haus8 siRNA (L-031247-01-0005; Dharmacon), 100 nM CENP-E siRNA (L-003252-000010; Dharmacon), 100 nM Kif4A siRNA (sc-60888; Santa Cruz Biotechnology), 100 nM Kif18A siRNA (4390825; Ambion), 100 nM Kid/Kif22 siRNA (4392420; Ambion), and 100 nM MKLP1 siRNA (sc-35936; Santa Cruz Biotechnology). After 4 h of incubation with transfection mixture, medium was replaced with regular cell culture medium. All experiments on siRNA-treated cells were perfomed 24 h after transfection, except for Haus8 siRNA-depleted cells, where silencing was done for 48 h. All treatments include at least three independent experiments.

### Speckle microscopy

Cells grown in glass coverslip dishes were stained with 1 nM SiR-tubulin dye (Spirochrome AG). After 15 min of staining, confocal live imaging was performed on a Dragonfly spinning disk confocal microscope system (Andor Technology) using 63x/1.47 HC PL APO glycerol objective (Leica) and Zyla 4.2P scientific complementary metal oxide semiconductor (sCMOS) camera (Andor Technology), and Expert Line easy3D STED microscope system (Abberior Instruments) using 60x/1.2 UPLSAPO 60XW water objective (Olympus) and avalanche photodiode (APD) detector. Images were acquired using Fusion software and Imspector software. During imaging, cells were maintained at 37°C and 5% CO2 within heating chamber (Okolab). For live imaging of RPE1 cells expressing CENP-A-GFP and centrin1-GFP, and stained with SiR-tubulin, 488-nm and 640-nm laser lines for Dragonfly microscope system, and 485-nm and 640-nm for Expert Line microscope system were used to excitate GFP, and SiR, respectively. In order to visualize SiR-tubulin speckles, images were acquired with 80% laser power and exposure of 1 s. Image acquisition was done on one focal plane every 5 or 10 s. Note that time-frame within which SiR-tubulin, at 1 nM concentration, can be visualized in patches on the mitotic spindle is between 15 and 75 min after SiR-tubulin staining.

### Immunostaining

Cells were fixed in ice-cold methanol for 1 min, except for astrin immunostaining experiment where cells were fixed in 37°C warm 4% paraformaldehyde for 10 min, and permeabilized for 15 min in 0.5% Triton X-100 in PBS. Following permeabilization, cells were blocked with 1% NGS in PBS for 1 h and incubated with primary antibodies at 4°C overnight. Primary antibodies were prepared in 1% NGS in PBS to 1:100 dilution. Following incubation with primary antibodies, cells were incubated with fluorescence-conjugated secondary antibodies at room temperature for 1 h. Secondary antibodies were prepared in 2% NGS in PBS to 1:250 dilution. To visualize DNA, cells were stained with DAPI for 10 min. After each step, cells were washed three times in PBS for 5 min. Primary antibodies used were: mouse anti-NuMA (sc-365532; Santa Cruz Biotechnology), mouse anti-PRC1 (sc-376983; Santa Cruz Biotechnology), rabbit anti-CENP-E (C7488; Sigma-Aldrich), mouse anti-Kif4A (sc-365144; Santa Cruz Biotechnology), rabbit anti-Kif18A (A301-080A; Bethyl Laboratories), mouse anti-Kid (sc-390640; Santa Cruz Biotechnology), rabbit anti-MKLP1 (ab174304, Abcam), and mouse anti-astrin (MABN2487; Sigma-Aldrich). Secondary antibodies used were: donkey anti-mouse IgG-Alexa 594 (Abcam), donkey anti-rabbit IgG-Alexa 594 (Abcam), and donkey anti-rabbit IgG-Alexa 647 (Abcam). Immunostained cells were imaged using Bruker Opterra Multipoint Scanning Confocal Microscope (Bruker Nano Surfaces) with a Nikon CFI Plan Apo VC 100x/1.4 numerical aperture oil objective (Nikon). 405/488/561/640-nm laser lights were used with following emission filters: BL HC 525/30, BL HC 600/37 and BL HC 673/11 (Semrock). Images were captured with an Evolve 512 Delta Electron Multiplying Charge Coupled Device (EMCCD) Camera (Photometrics) using a 200 ms exposure time.

### Image analysis

Measurements were performed in Fiji/ImageJ (National Institutes of Health). Quantification and data analysis were performed in R (R Foundation for Statistical Computing) and MATLAB (MathWorks). Figures and schemes were assembled in Adobe Illustrator CC (Adobe Systems). Statistical analysis was performed using Student’s t-test and Mann-Whitney test.

Upon inspection of tubulin speckle movement within the spindle, speckles which could be followed for at least 30 s were taken into account. For every tubulin speckle position, corresponding CENP-A and centrin positions, representing the location of sister kinetochores and spindle poles, respectively, were also tracked. Tracking was done by using the *Multi-point* tool. Speckles which started at a proximal kinetochore were categorized as a part of k-fiber, whilst speckles which started between sister kinetochores or proximal to the distal pole and passed through sister kinetochores were categorized as a part of bridging fiber. Speckles which could not be categorized as a part of k-fiber or bridging fiber were termed “other”. Speckle-pole velocity was calculated by fitting linear regression on distances between the tubulin speckle and the spindle pole towards which speckle is moving to during first 30 s of its trajectory.

For kinetochore alignment and inter-kinetochore distance measurements, the *Multipoint* tool was used to track positions of sister kinetochore pairs. The equatorial plane was defined with two points placed between outermost pairs of kinetochores on the opposite sides of the spindle. Kinetochore alignment was calculated as the distance between the midpoint of kinetochore pairs and the equatorial plane, while inter-kinetochore distance as the distance between two sister kinetochores.

In cells immunostained for PRC1, a 5-pixel-thick segmented line was used to track the pole-to-pole contour of PRC1-labeled overlap region. The pole-to-pole tracking was performed on single z-planes and the mean value of the cytoplasm was subtracted from the retrieved intensity profiles. The overlap length was determined as the width of the peak of the signal intensity in the central part of the contour in SciDavis (Free Software Foundation Inc.). The width of the peak was measured at the base of the PRC1 intensity peak where the PRC1 signal is roughly equal to the mean value of the PRC1 signal along the contour on either side of the peak.

To determine the percentage of protein depletion, we measured mean spindle intensity by encompassing the area of the spindle with the *Polygon selection* tool. Mean background intensity in the cytoplasm, measured using a 1x1 µm rectangle, was subtracted from the mean spindle intensity.

## ACKNOWLEDGEMENTS

We thank Alexey Khodjakov, Marin Barisic, Helder Maiato and Andrew McAinsh for the cell lines, Ivana Šarić for the drawings, and all members of Tolić and Pavin groups for helpful discussions. This work was funded by the European Research Council (ERC Synergy Grant, GA Number 855158, granted to I.M.T. and N.P., and ERC Consolidator Grant, GA Number 647077, granted to I.M.T.), the Croatian Science Foundation (HRZZ, project IP-2014-4753, granted to I.M.T.), and the QuantiXLie Center of Excellence, a project co-financed by the Croatian Government and European Union through the European Regional Development Fund—the Competitiveness and Cohesion Operational Programme (Grant KK.01.1.1.01.0004). The work of the doctoral students M.J. and A.B. was supported by the “Young researchers’ career development project – training of doctoral students” of the Croatian Science Foundation.

## AUTHOR CONTRIBUTIONS

P.R. and M.J. performed all experiments. P.R. analyzed the data with help of M.J. D.B. developed the model, based on A.B’s pilot work. N.P. and I.M.T. conceived the project and supervised the theory and experiments.

## SUPPLEMENTARY MATERIALS

**Figure S1.**
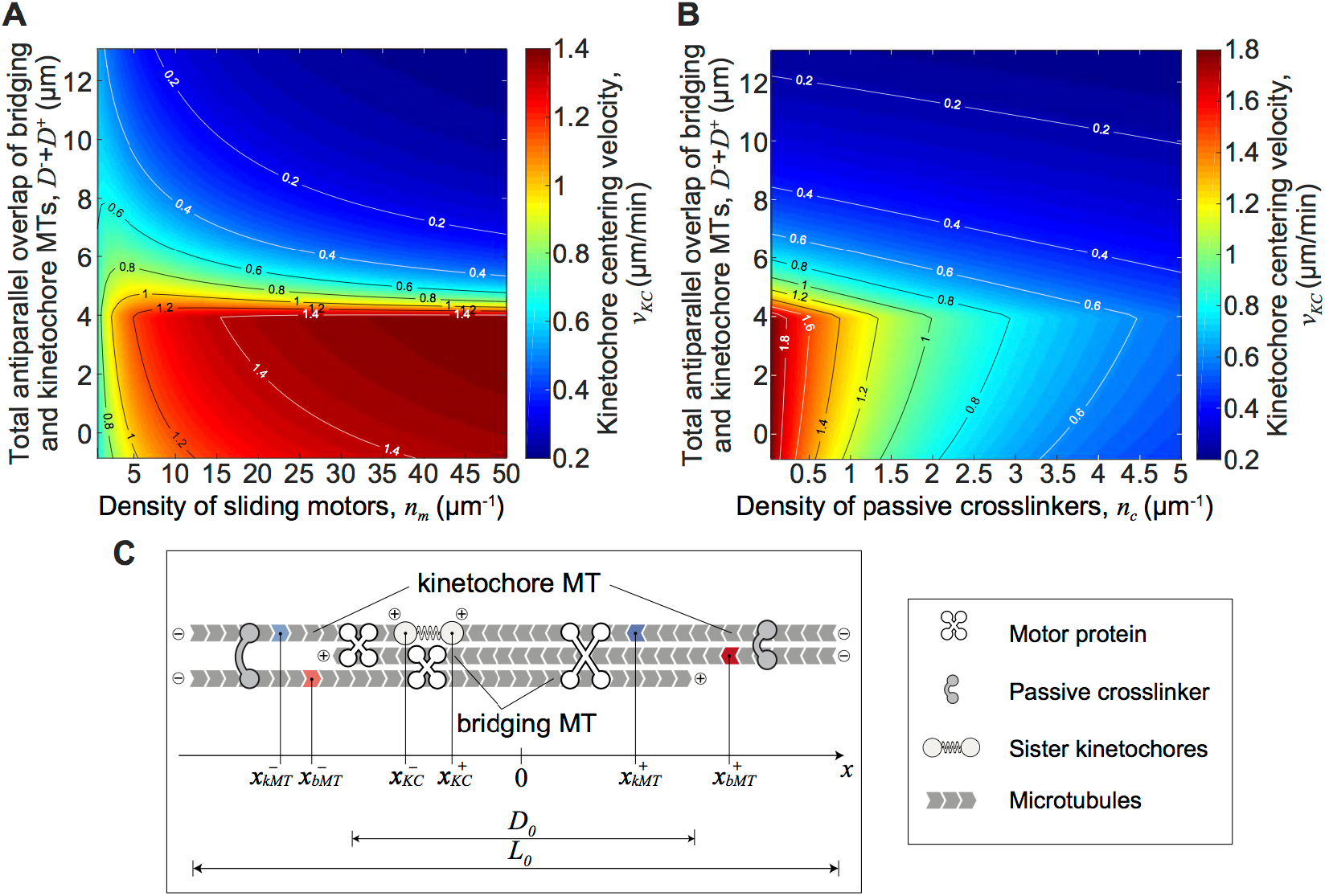
Kinetochore velocities for different parameters. (**A**) Velocity of kinetochore displaced 2 µm from center for different values of motor density and length of total antiparallel overlap of bridging and kinetochore microtubules. These data show two distinct regimes of centering velocity: Fast centering velocities for shorter overlaps (red region) which sharply decreases when the overlap length exceeds 4 µm (blue region). This abrupt change occurs when the shorter k-fiber loses connection with antiparallel bridging microtubule and thus there is no motor force that opposes centering movement. Transition between these two regimes is less abrupt for lower values of motor densities (*n*_m_ < 10 µm^-1^). (**B**) Velocity of kinetochore displaced 2 µm from center for different values of crosslinker density and length of total antiparallel overlap of bridging and kinetochore microtubules. Similar to panel (A), fast centering velocities are obtained for shorter overlaps (red region). However, the centering velocity decreases with the increase in crosslinker density, irrespective of the overlap length (blue region). Bars in (**A**) and (**B**) denote the relationship between color and velocity values. (**C**) Left: Scheme of the model. Kinetochore microtubules extend from the edges toward elastically connected kinetochores and bridging microtubules extend from the edges towards each other. Motor proteins connect antiparallel microtubules, while passive crosslinkers connect parallel microtubules. Total lengths of antiparallel and parallel microtubule overlaps are denoted as *D*_0_ and *L*_0_, respectively. Positions of sister kinetochores are marked on the x-axis. Positions of k-fibers (blue) and bridging fibers (red) are taken as arbitrary positions along their lattice and are also marked on the x-axis. Superscripts ***+*** and ***–*** denote the right and left sides, respectively. Right: Legend describing symbols for different elements of the spindle in the scheme. Parameters for all panels are given in **Fig. 1 B** if not stated otherwise.

**Figure S2.**
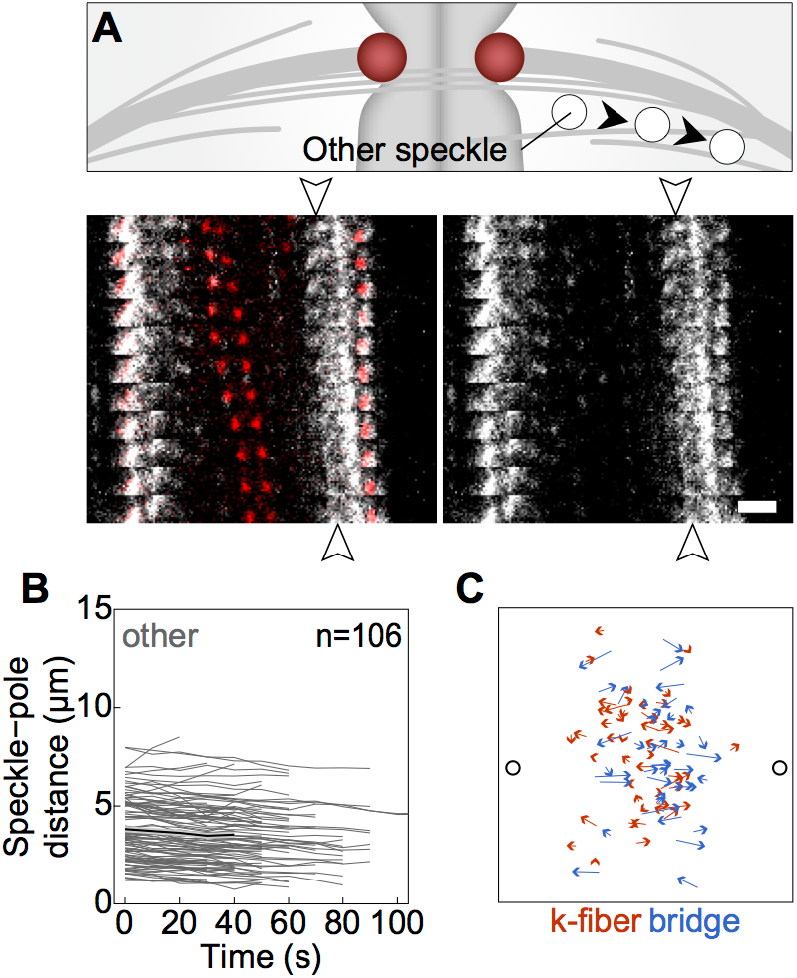
(**A**) Scheme of “other” speckles for which it could not be determined the type of microtubule they belong to (top). Montage over time demonstrating the movement of this group of speckles. Merge (left); tubulin channel only (right). Arrowheads mark starting and ending positions of the tracked speckle. Scale bar, 2 µm. (**B**) Distance of “other” speckles from the pole over time in untreated cells. Grey lines show individual speckles. Black line; mean. Grey area; s.e.m. (**C**) Trajectories of speckles belonging to k-fibers (red) and bridging fibers (blue) within 30 s of their movement. Arrows are pointing towards corresponding direction. Black circles; spindle poles.

**Figure S3.**
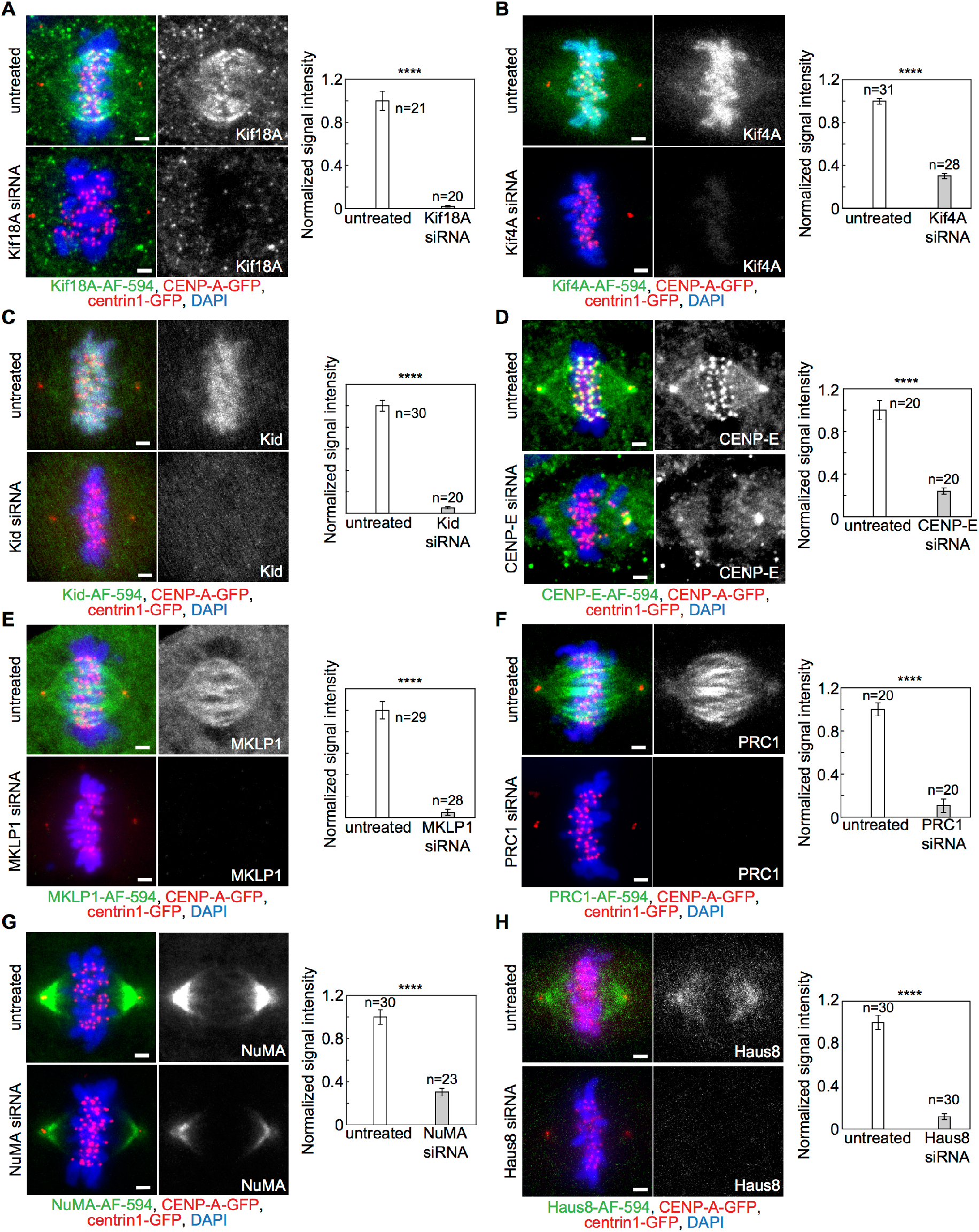
Fixed spindles in RPE1 cell line stably expressing CENP-A-GFP and centrin1-GFP (red) in cells immunostained for (**A**) Kif18A (AF-594, green), (**B**) Kif4A (AF-594, green), (**C**) Kid (AF-594, green), (**D**) CENP-E (AF-594, green), (**E**) MKLP1 (AF-594, green), (**F**) PRC1 (AF-594, green), (**G**) NuMA (AF-594, green) and (**H**) Haus8 (AF-594, green) in untreated (upper rows) and corresponding siRNA-treated cells (bottom rows), with DNA stained with DAPI (blue). Left: merge; right: protein of interest (grey). Graphs showing intensities of indicated proteins in untreated (white bars) and siRNA treated (grey bars) cells are given on the right. All values are normalized to the mean intensity value of untreated cells for each protein. All treatments include at least two independent experiments. n; number of cells. Scale bars; 2 μm. All images are maximum intensity projections of five z-planes smoothed with 0.5-pixel-sigma Gaussian blur. Statistical analysis, t-test. p-value legend as in **Fig. 3**.

**Figure S4.**
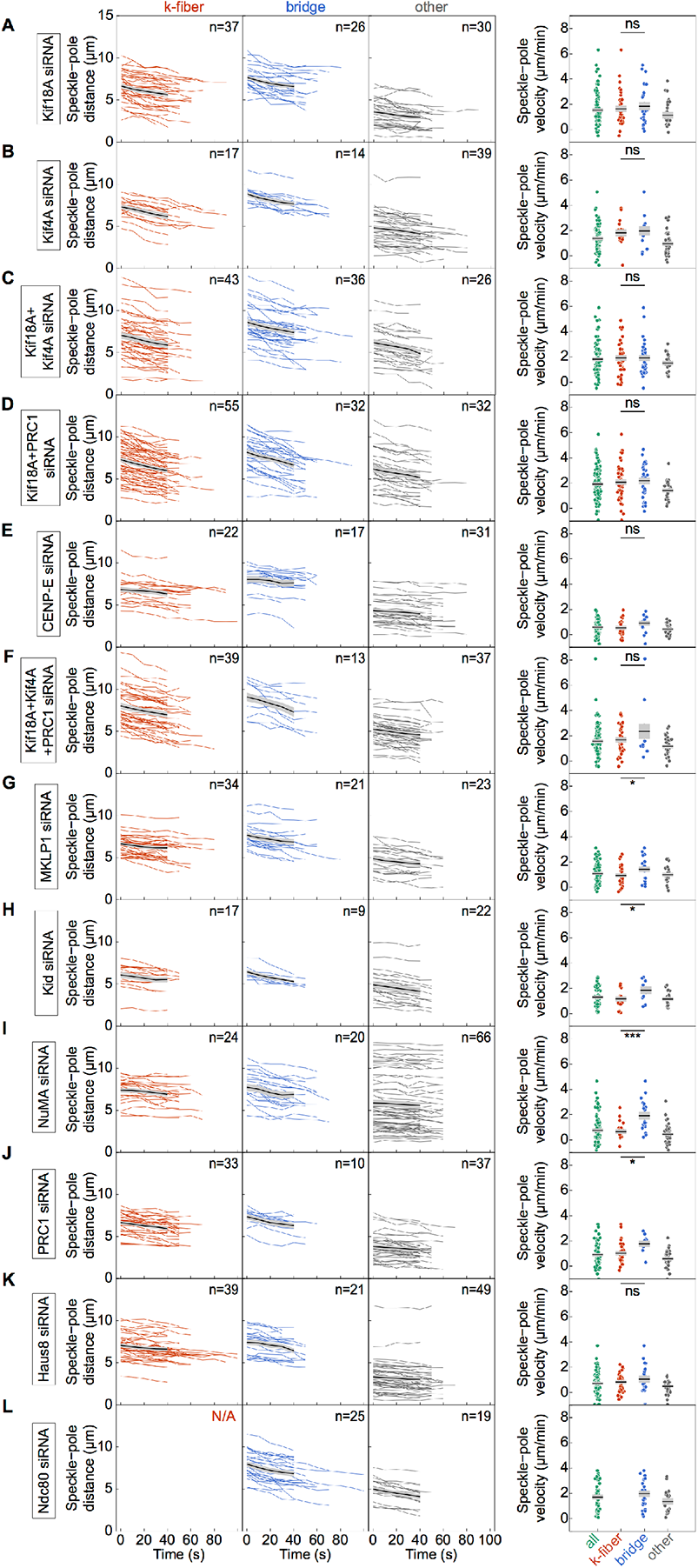
Poleward flux after depletion of (**A**) Kif18A, (**B**)Kif4A, (**C**) Kif18A and Kif4A, (**D**) Kif18A and PRC1, (**E**) CENP-E, (**F**) Kif18A, Kif4A and PRC1, (**G**) MKLP1, (**H**) Kid, (**I**) NuMA, (**J**) PRC1, (**K**) Haus8 and (**L**) Ndc80. Graphs from left to right show: speckles on kinetochore microtubules, speckles on bridging microtubules, and other speckles. Colored lines show individual speckles. Black line; mean. Grey area; s.e.m. Poleward velocity of the speckles is shown at the right. Black line; mean. Grey area; s.e.m. All treatments include at least three independent experiments. n; number of measurements. Statistical analysis, t-test. p-value legend as in **Fig. 3**.

**Figure S5.**
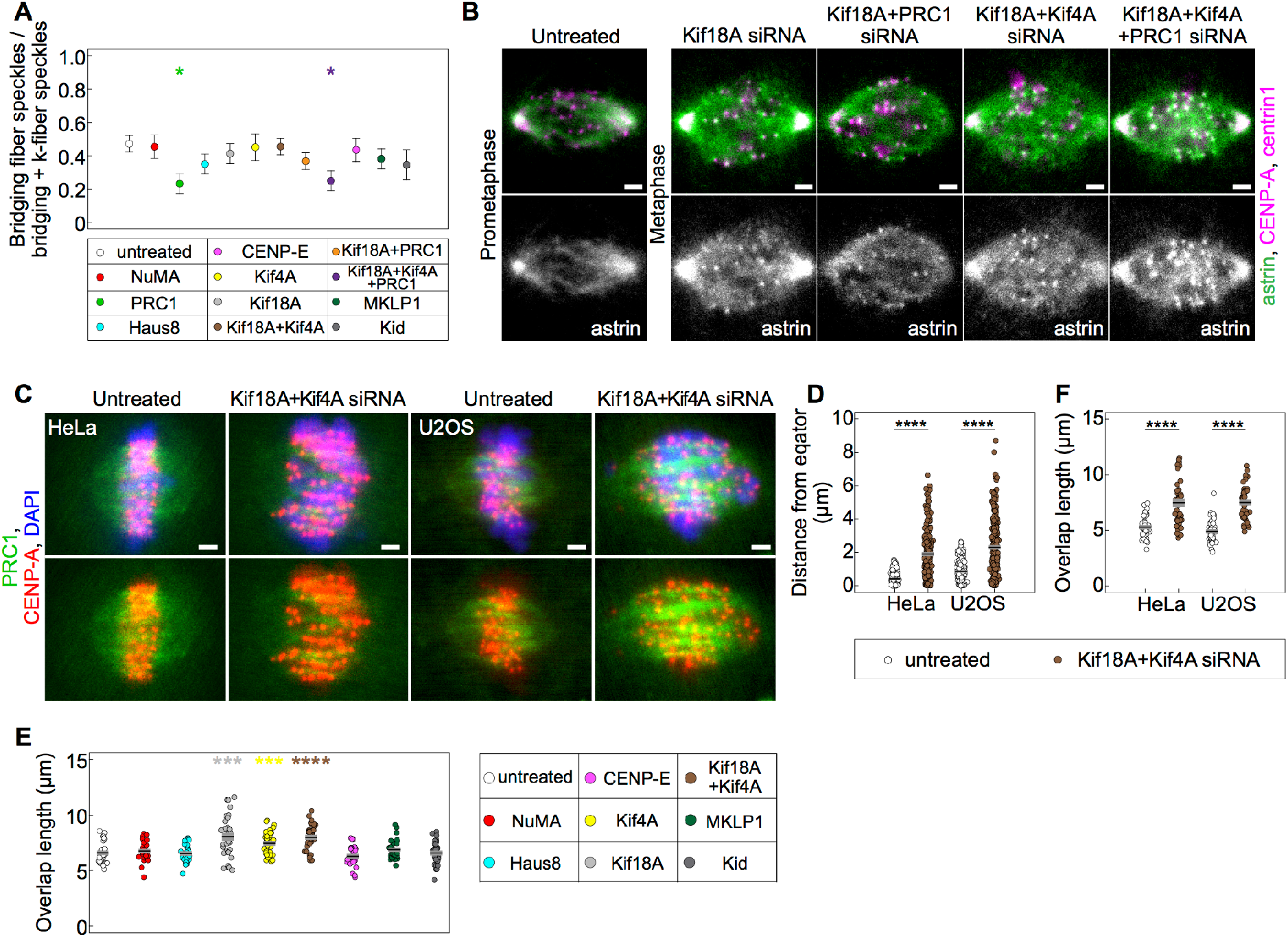
(**A**) Ratio of tracked speckles within bridging fibers and bridging and k-fibers (top) color-coded for corresponding treatments as in legend (bottom). Treatments include at least three independent experiments. (**B**) Fixed spindles in RPE1 cells stably expressing CENP-A-GFP and centrin1-GFP (magenta), immunostained for astrin (AF-594, green) in untreated and treated with Kif18A, Kif18A and PRC1, Kif18A and Kif4A, and Kif18A, Kif4A and PRC1 siRNA (left to right). Top: merge; bottom: astrin (grey). Images are sum intensity projections of five z-planes (**C**) Fixed spindles in HeLa and U2OS cells stably expressing CENP-A-GFP (red) in untreated (left) and Kif18A and Kif4A siRNA treated cells (right), immunostained for PRC1 (AF-594, green) and stained with DAPI (blue). Upper row; merge, bottom row; GFP and AF-594. Images are maximum intensity projections of five z-planes. (**D**) Kinetochore distance from equator in untreated and Kif18A and Kif4A siRNA treated HeLa (n = 172 and n = 235 kinetochore pairs) and U2OS (n = 216 and n = 281 kinetochore pairs) cells. (**E**) Length of PRC1-labeled overlaps. siRNA treatments are color-coded according to the legend (left). (**F**) Length of PRC1-labeled overlaps in untreated and Kif18A and Kif4A siRNA treated HeLa (n = 46 and n = 49 PRC1 bundles) and U2OS (n = 47 and n = 41 PRC1 bundles) cells. Black line; mean. Grey area; s.e.m. In (**A**, **E**), each treatment is compared with untreated cells. Statistical analysis, t-test, Mann-Whitney test (**D**). p-value legend as in **Fig. 3**.

**Movie 1.** Schematic movie showing numerical results of the theoretical model for chromosome alignment. Length of kinetochore microtubules (red), length of bridging microtubules (blue) and positions of elastically connected kinetochores (spring connecting circles) are obtained by numerical calculations. Several motor proteins (white X-shapes) are depicted as examples to visualize antiparallel overlap regions in which they exert forces. Kinetochores were initially displaced 3.3 μm, overlap length was *D*_>_ = 6.6 μm, and total simulation time was *t* = 250 s. The remaining parameters are given in **Fig. 1 B**.

**Movie 2.** A spindle in an untreated RPE1 cell stably expressing CENP-A-GFP and centrin1-GFP (red) and stained with SiR-tubulin dye (grey). Scale bar, 2 µm.

**Movie 3.** Spindles in CENP-E (left) and Ndc80 (right) siRNA treated RPE1 cells stably expressing CENP-A-GFP and centrin1-GFP (red) and stained with SiR-tubulin dye (grey). Scale bar, 2 µm.

**Movie 4.** Spindles in Kif18A (top left), Kif18A and Kif4A (top right), Kif18A and PRC1 (bottom left), and Kif18A, Kif4A and PRC1 (bottom right) siRNA treated RPE1 cells stably expressing CENP-A-GFP and centrin1-GFP (red) and stained with SiR-tubulin dye (grey). Scale bar, 2 µm.

**Movie 5.** A spindle in NuMa siRNA treated RPE1 cell stably expressing CENP-A-GFP and centrin1-GFP (red) and stained with SiR-tubulin dye (grey). Scale bar, 2 µm.

